# TopNEXt: Automatic DDA Exclusion Framework for Multi-Sample Mass Spectrometry Experiments

**DOI:** 10.1101/2023.02.16.527961

**Authors:** Ross McBride, Joe Wandy, Stefan Weidt, Simon Rogers, Vinny Davies, Rónán Daly, Kevin Bryson

**Affiliations:** School of Computing Science, University of Glasgow, Glasgow, United Kingdom; Glasgow Polyomics, University of Glasgow, Glasgow, United Kingdom; School of Mathematics and Statistics, University of Glasgow, Glasgow, United Kingdom

## Abstract

**Motivation:** Liquid Chromatography Tandem Mass Spectrometry (LC-MS/MS) experiments aim to produce high quality fragmentation spectra which can be used to identify metabolites. However, current Data-Dependent Acquisition (DDA) approaches may fail to collect spectra of sufficient quality and quantity for experimental outcomes, and extend poorly across multiple samples by failing to share information across samples or by requiring manual expert input.

**Results:** We present TopNEXt, a real-time scan prioritisation framework that improves data acquisition in multi-sample LC-MS/MS metabolomics experiments. TopNEXt extends traditional DDA exclusion methods across multiple samples by using a Region of Interest (RoI) and intensity-based scoring system. Through both simulated and lab experiments we show that methods incorporating these novel concepts acquire fragmentation spectra for an additional 10% of our set of target peaks and with an additional 20% of acquisition intensity. By increasing the quality and quantity of fragmentation spectra, TopNEXt can help improve metabolite identification with a potential impact across a variety of experimental contexts.

**Availability:** TopNEXt is implemented as part of the ViMMS framework and the latest version can be found at https://github.com/glasgowcompbio/vimms. A stable version used to produce our results can be found at 10.5281/zenodo.7468914. Data can be found at 10.5525/gla.researchdata.1382.

**Supplementary information:** Supplementary data are available at *Bioarxiv* online.

## 1 Introduction

Liquid chromatography (LC) tandem mass spectrometry (MS/MS) is commonly used to identify small molecules in untargeted metabolomics. On its own a mass spectrometer may identify the relative abundances (intensities) of different masses of ion (*m/z*, mass-to-charge ratio) of an injected sample. The use of liquid chromatography coupled to electrospray ionisation (ESI) generates ions into the mass spectrometer separated in time, creating three-dimensional data in which intensity profiles of different ions (chromatographic peaks) may be observed across retention time (RT). In MS/MS schemes, we may isolate ions within a fixed mass range (an isolation window), fragment them, and measure the intensities and *m/z* of the fragments. Therefore LC-MS/MS combines MS1 (survey) scans, which report intensities of all ions currently eluting from the chromatographic column, and MS2 (fragmentation) scans, which each produce measurements of the intensities of fragment ions — a fragmentation spectrum. A combination of MS1 and MS2 scans will produce the data which allows us to annotate chemicals by directly matching their fragmentation spectra to spectral databases, by machine-learning assisted comparison with structural databases (Dührkop *et al*., 2019; Djoumbou-Feunang *et al*., 2019) or by analysis with metabolome data-mining tools (Wang *et al*., 2016; van Der Hooft *et al*., 2016). However, biological samples are often highly complex and may contain hundreds or thousands of metabolites. Consequently, we must make a careful choice of fragmentation strategy to decide assignments of MS1 and MS2 scans — a well-designed fragmentation strategy should produce as many relevant fragmentation spectra as possible, at the highest quality possible.

DDA (Data-Dependent Acquisition) methods are often observed to produce a lower number of spectra compared to DIA (Data-Independent Acquisition) methods (Guo and Huan, 2020; Wandy *et al*., 2023). DIA methods set in advance a scan schedule in which MS2 scans fragment all ions within a large isolation window, whereas DDA methods use real-time feedback from MS1 scans to target a single precursor ion at a time. Consequently, DIA scans require additional processing to separate the hybrid spectra produced and algorithms for this purpose are an area of ongoing research (Tsugawa *et al*., 2015; Tada *et al*., 2020). DDA MS2 scans typically produce higher quality, ready-to-use spectra, but are instead bounded by their ability to optimally schedule scans for any given sample. Previous work (Davies *et al*., 2021) has shown that it is theoretically possible to extract many more spectra from a given sample with the correct DDA schedule, so in this work we will focus on improving DDA scan-scheduling in practice. Currently, the most widely used DDA method is TopN, which repeatedly schedules a duty cycle of one MS1 scan followed by up to *N* MS2 scans. In each of these MS2 scans one of the top *N* most intense precursor ions observed in the last MS1 scan is targeted, and as a consequence TopN often wastes time repeatedly recollecting spectra for the most abundant precursor ions rather than collecting new spectra. To counteract this effect, TopN data is normally acquired using Dynamic Exclusion Windows (DEWs), which are a (*rt, m/z*) box around each fragmentation event, forbidding further fragmentation events to fall within a DEW’s *m/z* tolerance until a specified period of time has elapsed. More recent extensions of this idea include SmartRoI and WeightedDEW (Davies *et al*., 2021) which add more flexible criteria for allowing refragmentation.

In multi-injection, multi-sample experiments, several samples are injected in series. If the same high intensity traces appear across multiple injections, a naive TopN will repeatedly fragment the same molecular ions. Therefore TopN is often augmented with static exclusion windows which may persist across injections and, like the DEW, forbid fragmentation within their bounds. An obvious approach to these iterative exclusion schemes is to remember DEWs between samples for further use as exclusion windows (Bendall *et al*., 2009). However exclusion lists are instead frequently created from manual analysis and existing software tools are generally only semi-automated (Koelmel *et al*., 2017). A contrasting approach is to analyse samples offline and pre-schedule scans targeting individual ions (Broeckling *et al*., 2018; Zuo *et al*., 2021). However, in a multi-sample context this approach may have difficulty when its plan differs from reality, due to random variation between injections or genuine biological variation between samples (Wandy *et al*., 2019). The Thermo vendor method AcquireX (Thermo Fisher Scientific, 2020) combines offline processing with real-time DDA decision-making, but can only be used to process repeated injections of the same sample.

We introduce TopNEXt, a real-time DDA scan-prioritisation framework and an extension of our previously introduced Virtual Metabolomics Mass Spectrometer (ViMMS) (Wandy *et al*., 2022). TopNEXt implements several improved multi-sample fragmentation strategies within a modular and cohesive base. To do this, it extends the concept of exclusion windows by implementing novel ideas of intensity exclusion (where we may revisit high intensity signals) and Region of Interest (RoI) area exclusion (where we compare entire groups of MS1 points for similarity against exclusion windows) in addition to existing concepts of real-time RoI-tracking and multi-sample exclusion (Davies *et al*., 2021; Bendall *et al*., 2009). We show through both simulated and lab experiments that these concepts enable collecting more and higher-quality relevant fragmentation spectra compared to TopN, allowing DDA strategies to obtain more metabolite annotations in future.

## 2 Methodology

TopNEXt is embedded within the open-source Python-based ViMMS (Wandy *et al*., 2022) framework. ViMMS allows us to implement new fragmentation strategies in Python and test their performance using either re-simulated data or an actual mass spectrometer — currently ViMMS can control only Thermo Fisher IAPI instruments (Thermo Fisher Scientific, 2016). In either case, we can evaluate these fragmentation .mzMLs (Martens *et al*., 2011) against an aligned peaklist produced from corresponding fullscan .mzMLs via peak-picking with e.g. MZMine 2 (Pluskal *et al*., 2010). We use the metrics of *peak coverage*, a measure of how many detected chromatographic peaks we have collected fragmentation spectra for, and *intensity coverage*, a measure of the intensity at which we collected spectra for detected peaks (a proxy for quality).

For our experiments we collected ten different store-bought beers and ran them in four batches on four separate days. The first batch was used to optimise our fragmentation strategy parameters, and the other three to produce data for our experiments. Beer was chosen because it is complex and chemically diverse but is also easy to obtain. Sample extraction was done by adding chloroform and methanol in a 1:1:3 ratio (v/v/v) and mixing with a vortex mixer. The extracted solution was then centrifuged to remove protein and other precipitates, and the supernatant was stored at -80°C. Chromatographic separation with HILIC was performed on all samples by injecting 10 *µ*L beer extract with a Thermo Scientific UltiMate 3000 RSLC liquid chromatography system and a SeQuant ZIC-pHILIC column. A gradient elution was carried out with 20 mM ammonium carbonate (A) and acetonitrile (B), starting at 80% (B) and ending at 20% (B) over a 15 min period, followed by a 2 min wash at 5% (B) and a 9 min re-equilibration at 80% (B). The flow rate was 300 *µ*L/min and the column oven temperature was 40°C. Mass spectra data was generated using a Thermo Orbitrap Fusion tribrid-series mass spectrometer controlled by Thermo IAPI via ViMMS. Full-scan spectra were acquired in positive mode with a resolution of 120,000 and a mass range of 70-1000 *m/z*. Fragmentation spectra were acquired using the orbitrap mass analyzer at a resolution of 7,500, with precursor ions isolated using a 0.7 *m/z* width and fragmented using a fixed HCD collision energy of 25%. The ACG was set at 200,000 for MS1 scans and 30,000 for MS2 scans.

The exact beers used, the parameters used for peak-picking, thorough descriptions of our evaluation metrics and the specifics of our controller parameter optimisation procedure can be found in Supplementary Sections 1, 2, 3 and 4.

## 3 Algorithms

In ViMMS, and therefore TopNEXt, DDA fragmentation strategies are expressed as a modular scoring function. In all the strategies we will discuss here, their scoring function is used to rank the precursors in each MS1 scan, so that up to *N* MS2 scans can be scheduled on the *N* most highly-scored precursors. TopN can be represented using the scoring function in Equation 1.

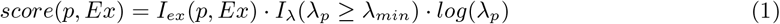

For TopN, all precursors *p* are scored directly by their log intensity *log*(*λ*_*p*_). To ensure that all acquisitions are of a usable quality, it is common practice to only consider precursors above a minimum intensity threshold *λ*_*min*_, where *I*_*λ*_(*λ*_*p*_ *≥ λ*_*min*_) is an indicator function expressing this constraint. We also implement the DEW via the exclusion indicator function *I*_*ex*_(*p, Ex*) which is 0 if the precursor *p* falls within any exclusion window in the set *Ex*, and 1 otherwise. If there are insufficient targets with a score above zero, a duty cycle may end with fewer than *N* MS2 scans.

TopNEXt implements several different strategies containing a number of features and these can be seen in Table 3. Because each strategy is built on TopN they are therefore implemented by using only the three terms in Equation 1 or more complicated expressions substituted in their place. For example, TopN is commonly extended to multi-sample contexts with additional exclusion windows which function identically to the DEW. For our fully automatic **TopNEX (TopN EXclusion)** implementation, we use an iterative exclusion scheme where DEW from previous injections are carried forward to the current injection as in (Bendall *et al*., 2009) — so we just include these additional exclusion windows in *Ex*. Sections 3.1-3.3 dissect the features and controllers in Table 3, showing how we can define them by substituting “modified intensities” in place of *λ*_*p*_. TopNEXt facilitates these operations efficiently with a simple geometry of points, lines and rectangles for which details can be found in Supplementary Section 6.2. Note that in our experiments later we will also substitute the SmartRoI and WeightedDEW weights (Davies *et al*., 2021) in place of *I*_*ex*_: this procedure is described in Supplementary Section 6.1.

### 3.1 Multi-Sample RoI Exclusion

RoIs are rectangular regions in *rt, m/z* space which are typically constructed as a first step in peakpicking, to group individual MS1 points along rt into approximately peak-like objects. By using ViMMS’ existing implementation of real-time RoI tracking (Davies *et al*., 2021) implemented with the centwave RoI-building algorithm (Tautenhahn *et al*., 2008) we can build and manipulate peak-like objects in place of fixed-size exclusion regions. In this scheme, we replace the standard DEW with the rule that no RoI can have another fragmentation event fall inside it within some retention time tolerance of its last fragmentation. To denote this change we substitute all instances of a precursor *p* with its containing RoI, *r*. Then *I*_*ex*_ assumes responsibility for this rule in addition to behaving as before, where for each RoI we check whether its precursor in the current MS1 scan falls into an exclusion window in the set *Ex*. To demonstrate that this alone does not significantly alter results we define **TopN RoI** which has an empty *Ex* (i.e. only RoIs are given as argument to *I*_*ex*_) and **TopN Exclusion RoI** with an *Ex* containing the DEWs that would have appeared in previous injections in a non-RoI method. The first new concept implemented by TopNEXt is to populate *Ex* not with remembered DEW boxes, but instead with exclusion windows matching RoIs fragmented in previous injections. In doing so, **Hard RoI Exclusion** extends RoI-tracking to between-injections exclusion also. Because we are only changing the contents of *Ex*, all three of these controllers can be expressed by Equation 2. An example of how Multi-Sample RoI Exclusion works can be seen in Figure 1: some points in the second injection fall within the area labelled *ab* and hence inside *a*, so under Hard RoI Exclusion would not be considered for fragmentation.

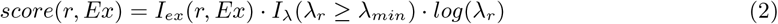

**Table 1:** A breakdown of which fragmentation strategies incorporate which features. The column “RoI DEW” describes whether the within-sample exclusion is tied to RoIs, whereas “Multi-Sample RoI Exclusion” (Section 3.1) shows whether between-sample exclusion is tied to RoIs. “Multi-Sample” indicates whether it carries over information between samples and “Intensity Exclusion” (Section (3.2)) and “RoI Area Weighting” (Section (3.3)) show whether the between-sample exclusion uses intensity changes or RoI area, respectively. All methods below the line break are implemented using the TopNEXt framework: those above are implemented elsewhere in ViMMS.

| Method | Multi-Sample | RoI DEW | Multi-Sample RoI Exclusion | Intensity Exclusion | RoI Area Weighting |
| --- | --- | --- | --- | --- | --- |
| TopN |  |  |  |  |  |
| TopN Exclusion | ✓ |  |  |  |  |
| TopN RoI |  | ✓ |  |  |  |
| TopN Exclusion RoI | ✓ | ✓ |  |  |  |
| Hard RoI Exclusion | ✓ | ✓ | ✓ |  |  |
| Intensity RoI Exclusion | ✓ | ✓ | ✓ | ✓ |  |
| Non-Overlap | ✓ | ✓ | ✓ |  | ✓ |
| Intensity Non-Overlap | ✓ | ✓ | ✓ | ✓ | ✓ |

**Figure 1:**
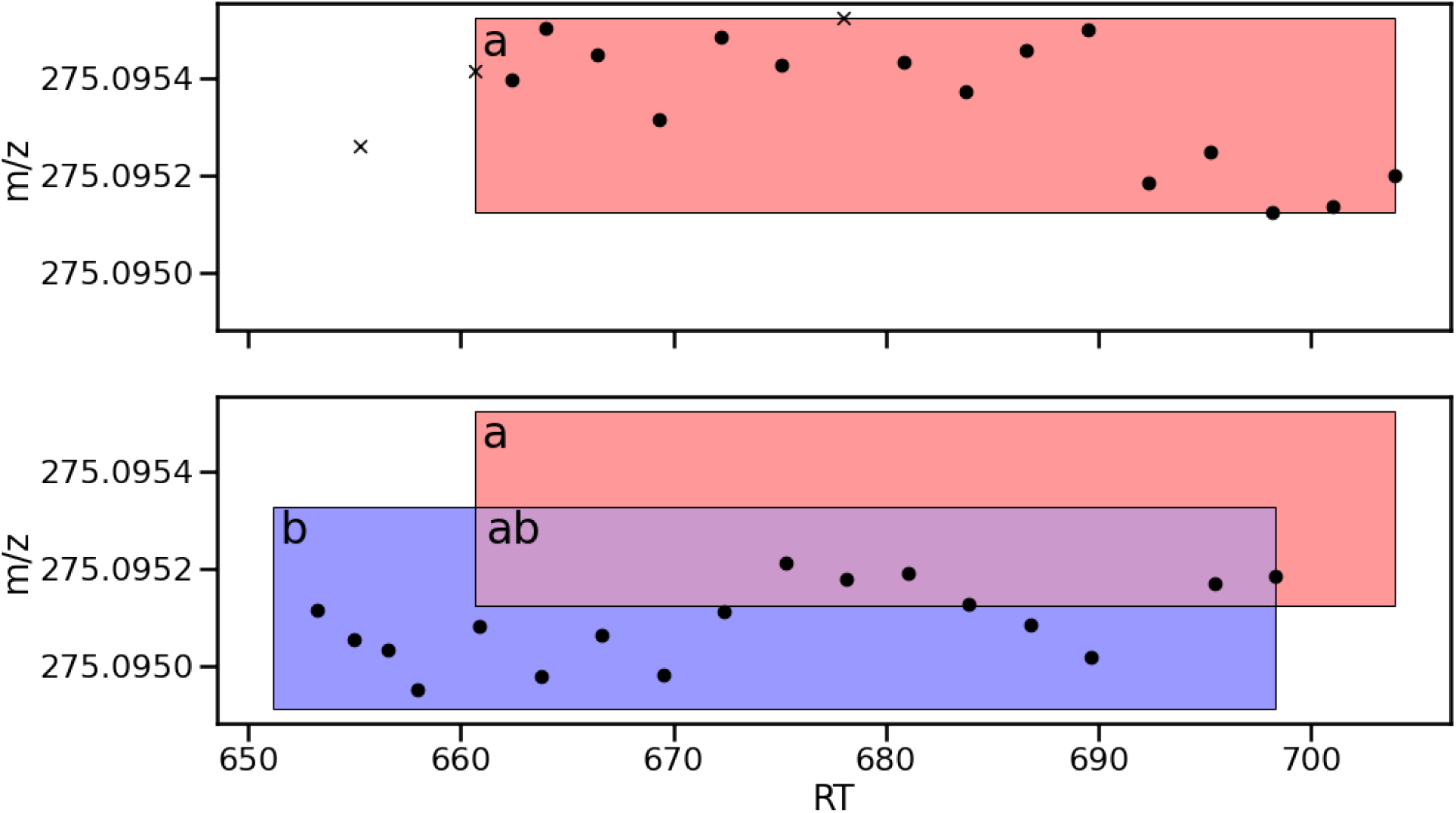
An illustration of RoI-tracking when using RoIs as exclusion windows, where from top to bottom each subplot represents a successive injection. The points are individual observations in MS1 scans. A cross represents the precursor of a fragmentation event. On the first injection, the RoI *a* is drawn. On the second injection, *a* persists as an exclusion window, while *b* is drawn around the new points, forming the overlapping area *ab*. Note that *a* and *b* are drawn here after all points were observed, but as RoIs would be dynamically extended to the right to cover the points as we observed them in real-time.

### 3.2 Intensity Exclusion

While exclusion regions by design prevent revisiting an area of the space, it may sometimes be desirable to do so. For example, if the fragmentation strategy has run out of opportunities to increase coverage, it may be desirable to reacquire a peak at a higher intensity, potentially improving spectral quality. To encourage this behaviour we replace the intensity *log*(*λ*_*r*_) in Equation 2 with a modified intensity value, where we reduce the current intensity of a ROI *r* by the highest intensity of *r* at any previous fragmentation. This is *log*(*λ*_*r*_) *− log*(*ϕ*(*r, Ex*)) where *ϕ*(*r, Ex*) is a function that computes the maximum intensity of any Multi-Sample RoI Exclusion windows the precursor falls within. *log*(*ϕ*(*r, Ex*)) is 0 if *r* has not been previously fragmented i.e. the intensity used in the score is not modified. This defines **Intensity RoI Exclusion**, which is shown in Equation 3. Note that *I*_*ex*_ only considers DEWs (which only exist within a single injection) and *ϕ*(*r, Ex*) only considers exclusion windows which persist across injections.

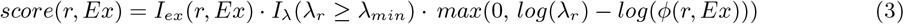

### 3.3 RoI Area Weighting

Hard RoI Exclusion and Intensity RoI Exclusion build on TopN Exclusion by checking whether a query RoI has its last precursor contained within previously fragmented RoIs. Instead we can compare entire RoIs for similarity: if the area of a query RoI is largely uncovered by previously fragmented RoIs, then it is likely it is a RoI we have not previously fragmented. Therefore TopNEXt implements a **Non-Overlap** strategy, which weights the intensity of the query RoI by how much of its area is uncovered by fragmentation boxes. When two rectangles (RoIs or exclusion windows) *a* and *b* overlap, they can be separated into a rectangular area where both *a* and *b* are present, and two non-rectangular areas where only one of *a* or *b* is present. It is possible to replace all three of these areas with an equivalent set of non-overlapping rectangles, and repeating this procedure will allow dissection of any set of overlapping rectangles into non-overlapping rectangles. We can then easily compute the area of the region where only *a* is present — *a*’s area of non-overlap — by summing the areas of the rectangles covering this region. To compute the Non-Overlap score we find this area for the query RoI as a proportion of its total area and use it as a [0, 1] bounded weight on the log intensity i.e. a power on the raw intensity by log laws. *λ*_*r*_ is therefore substituted by the modified intensity *prop*(*r*) · *log*(*λ*_*r*_), where *prop*(*a*) is the proportional area of non-overlap for *a*, defined in Equation 4. Here *a*_*i*_ is the *i*th rectangle covering *a*’s area of non-overlap, and 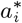 is the *i*th rectangle covering *a*.

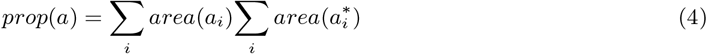

Equation 5 expresses the whole Non-Overlap scoring function.

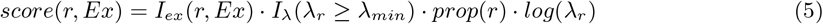

Then when deciding whether a point in RoI *b* should be fragmented in Figure 1, we would raise the intensity of the point *λ*_*r*_ to the power of the blue area *b* as a proportion of the total area of the RoI *b*.

**Intensity Non-Overlap** combines all the concepts introduced in Sections 3.1 - 3.3 i.e. we combine Non-Overlap with intensity scoring. As in Non-Overlap, the set of overlapping rectangles is split into non-overlapping rectangles. But while Non-Overlap uses only the rectangles which would overlap the query RoI, but not an exclusion window, Intensity Non-Overlap uses all rectangles which would overlap the query RoI. Firstly, each exclusion window has intensity equal to precursor intensity of its associated fragmentation event, and each RoI has intensity equal to its intensity in the most recent MS1 scan. Supposing that we are interested in calculating the Intensity Non-Overlap score for *a*, then unique combinations of overlapping rectangles including *a* can be written as *aB* where *B* ∈ {*ϵ, b, c, bc, d, bd, bc, bcd*…} for any overlapping boxes *b, c, d*… and with *ϵ* representing no other boxes. Each of these rectangles has a modified intensity associated with it equal to the difference between *a*’s intensity and the maximum intensity of any overlapping boxes, i.e. *λ*_*aB*_ = *λ*_*a*_ *− b*′ *∈ Bmax*(*λ*_*b*_*′*), where when *B* = *ϵ* we have *b*′ *∈ Bmax*(*λ*_*b*_*′*) = 0.

In Non-Overlap we only considered 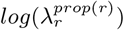. For Intensity Non-Overlap, we generalise this to the logarithm of the sum of all modified intensities given to each unique combination of boxes taken tothe power of their proportional area — this is shown in full in Equations 6 and 7.

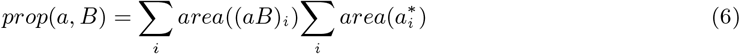

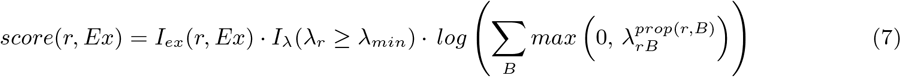

Then when deciding whether to fragment *b* in Figure 1, *b* and *ab* would have their intensity calculated as *λ*_*b*_ and *λ*_*b*_ *− λ*_*a*_ respectively, and their area as a proportion of the total area of *b* would be used as a power on this before finally summing them together. A detailed example of how Intensity Non-Overlap is computed can be found in Supplementary Section 6.3.

## 4 Results

We test our results on two scenarios — repeated injections of the same individual beer sample (multi-injection, single-sample), and injections of different beer samples, with repeats (multi-sample). In the first case, roughly the same peaks are encountered each time at the same *m/z* and RT position in an injection, which should cause it to be relatively predictable and therefore straightforward. The second case should have samples with partial overlap in metabolites, making optimally fragmenting peaks in the correct sample more challenging. Primarily we should expect that in the single-sample case especially TopN’s coverage will not significantly increase, but the coverage of multi-sample methods will, and we should expect that intensity-based methods will obtain more intensity coverage and continue to increase in intensity coverage even when coverage stops improving.

Section 4.1 contains simulated results combining all strategies built on top of TopNEXt with the three DEW variants (regular DEW, SmartRoI, WeightedDEW). We also present TopN, TopN RoI and TopN Exclusion as baselines. To differentiate the non-RoI implementation of TopN Exclusion with the RoI-based implementation within TopNEXt we denote them “TopN Exclusion” and “TopNEX” respectively. These simulated results were produced using the fullscans from our fourth batch of lab experiments, and MS1 and MS2 scan lengths were fixed to be 0.59 and 0.19 respectively — the average times from the same instrument in (Davies *et al*., 2021) — so they could be more exactly reproducible.

Section 4.2 contains the lab experiments. These have a significant instrument time cost to run, so for the multi-injection experiment (the third batch of our experiments) we only present comparison of TopN Exclusion, Non-Overlap and Intensity Non-Overlap. These three were chosen to compare performance of an intensity method to a non-intensity TopNEXt method and a baseline method as sample coverage becomes exhaustive. For the multi-sample experiment we tested all the main variants — TopN Exclusion, Non-Overlap and Intensity Non-Overlap in the second batch, and TopN, Hard RoI Exclusion and Intensity RoI Exclusion in the fourth. This shows performance on a complex and realistic scenario. In the lab experiments, all TopNEXt methods use WeightedDEW exclusion. WeightedDEW was found to have the best performance when optimising parameters on all three in simulation (see Supplementary Section 4).

### 4.1 Simulated Results - Resimulated Chemicals

#### 4.1.1 Multi-injection, Single Sample Results

In our simulated multi-injection results given in Figure 2A we have twenty injections of the same beer. As we expect, TopN is a completely flat line which does not improve beyond seeing the same sample once as no RT noise was introduced during simulation. The other controllers are all roughly competitive on coverage, with the gap being at most around 2% between the best and worst performing variants. The best performing variants are the different implementations of TopN Exclusion, and after ten samples most methods have converged to near-complete coverage of the sample. Despite gaining coverage the fastest, in intensity coverage the TopN exclusion variants perform the worst by a significant margin, which increases up to around 3% behind the worst new TopNEXt-based method, Hard RoI Exclusion. Intensity RoI Exclusion has significantly better intensity coverage than any non-intensity method and Intensity Non-Overlap is again better than Intensity RoI Exclusion by a significant margin, performing the best on this metric. For most methods in this example, but particularly the intensity methods, the SmartRoI variants are especially effective, with Intensity RoI Exclusion being approximately 6% higher in intensity coverage compared to Non-Overlap, and Intensity Non-Overlap being approximately 5% beyond that. In total Intensity Non-Overlap has an intensity coverage of 92%, a nearly 20% difference ahead of the nearest TopN Exclusion variant. Importantly, we can also observe that the TopN exclusion variants plateaus in intensity coverage shortly after doing so in coverage, and that intensity methods especially maintain a significant slope even as their coverage does not, plateauing later in the process.

**Figure 2:**
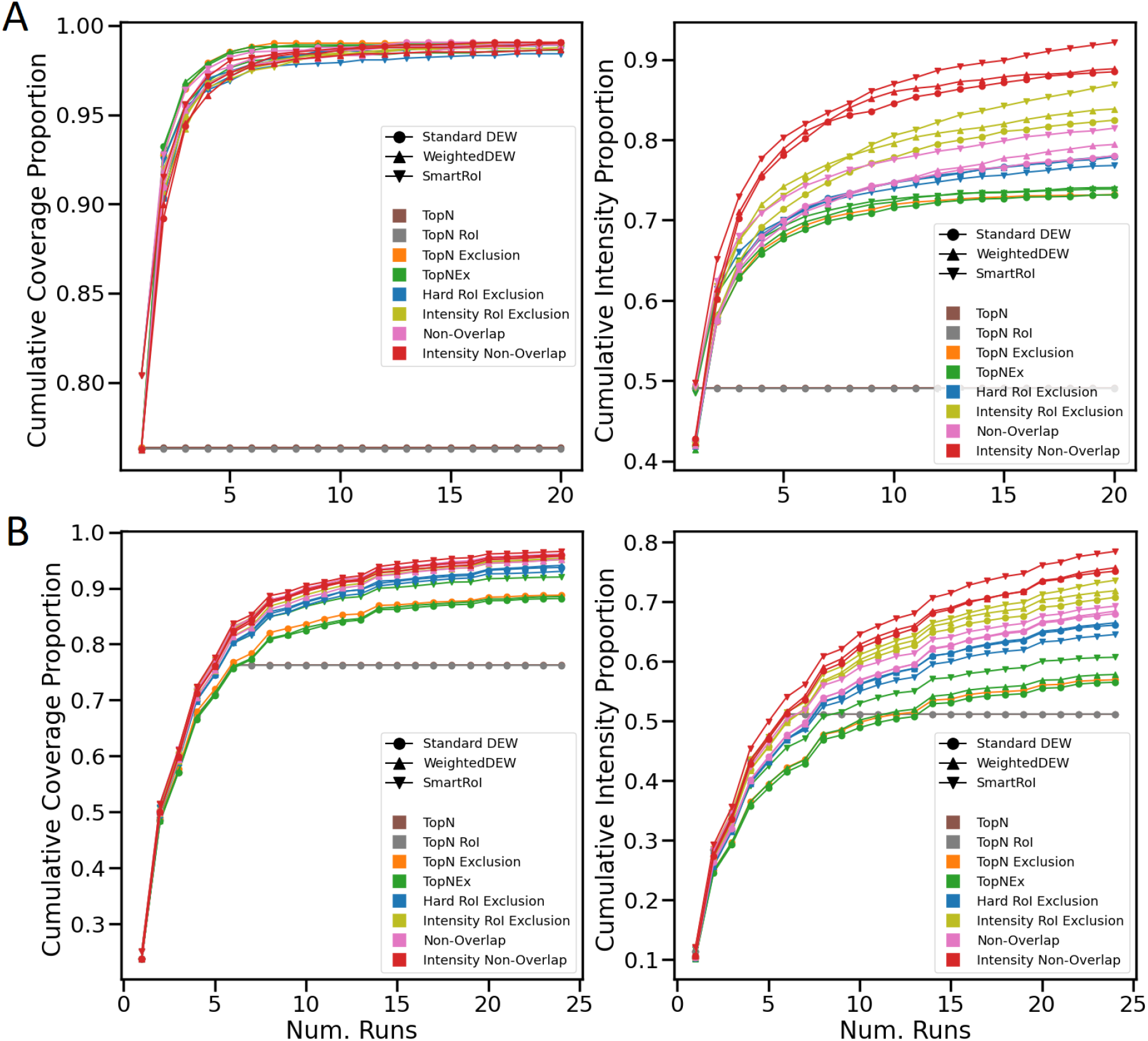
A: simulated experiment with the same beer repeated for twenty injections. B: simulated experiment with six different beers each repeated four times.

#### 4.1.2 Multi-Sample Results

For the multi-sample experiment we repeat six unique beers (labelled 1-6) four times each for each controller, in the order 1-2-3-4-5-6-1-2-3-4… meaning at any point no beer sample has been repeated more than once more than another. This should allow the strategies to firstly collect all shared metabolites, then collect those exclusive to some samples later, rather than potentially missing them permanently. When this ordering is applied to the lab experiment, it has the potential of causing experimental issues on the real instrument (e.g. through retention time drift) but we decided the benefit of additional collection opportunities outweighed the risk. Figure 2B shows the results of this 6-4 (6 samples, 4 repeats) experiment. Once again, TopN stops gaining any coverage or intensity coverage once new beers cease to be introduced, and the multi-sample methods all have a significant advantage over it. However, the methods rank differently in coverage this time, and while all new methods are roughly competitive in this respect, most of the TopN exclusion variants trail by a significant margin of around 4% behind the least effective of these methods, Hard RoI Exclusion. TopNEX SmartRoI is very close to Hard RoI Exclusion, but it still nonetheless ranks below all of the new methods. The intensity methods perform *best* on coverage here, with Intensity Non-Overlap being the best controller overall, with around 8% increase in coverage from the baseline TopN exclusion implementation to Intensity Non-Overlap SmartRoI. The differences in intensity coverage remain mostly similar to the same beer experiment, with Intensity Non-Overlap SmartRoI being roughly 21% ahead of baseline TopN Exclusion.

### 4.2 Lab Results

#### 4.2.1 Multi-injection, Single Sample Results

In the prior simulated results given in Section 4.1 coverage was often exhausted significantly before 10 injections, so we ran only 10 injections for the multi-injection experiment on the actual instrument. Figure 3A shows that all the multi-sample methods are competitive on coverage, with Intensity Non-Overlap being the lowest by a slight margin (as it is focusing on reacquiring peaks at higher intensities, i.e. intensity coverage). Both overlap methods have significantly better intensity coverage (with Intensity Non-Overlap having a further advantage) — but most notably it can be observed that the curves of the other two methods flatten as they run out of new peaks to acquire, but the Intensity Non-Overlap curve flattens at a decreased rate. This demonstrates the advantage it has in continuing to reacquire peaks at higher intensities even once coverage gains cease.

**Figure 3:**
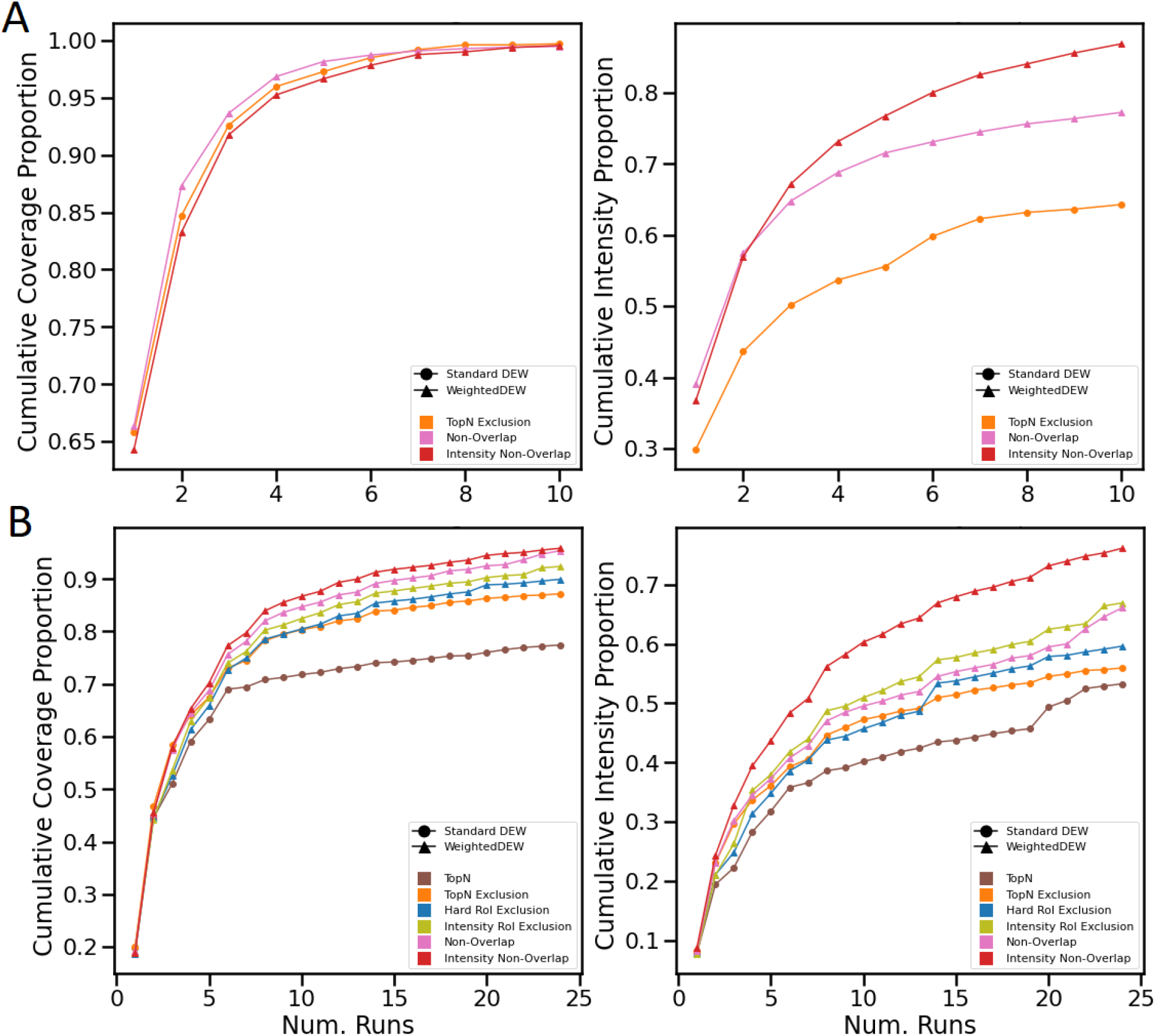
A: lab experiment with the same beer repeated for ten injections. B: lab experiment with six different beers each repeated four times.

#### 4.2.2 Multi-Sample Results

The 6-4 experiment was the most complex and representative of real-use, so we ran this again without changing the setup of runs or beers. Figure 3B shows that in both coverage and intensity coverage TopN is again the weakest of the methods and TopN exclusion trails behind the new TopNEXt methods. The new methods are competitive in terms of coverage: the “intensity” methods clearly improve the intensity coverage by a large margin. There are some particularly large spikes in intensity coverage (especially around sample 20) but the overall trend, however, reassuringly matches the simulated results. To further support these results, Supplementary Section 5 contains a number of other simulated experiments showing the generalisation performance of these methods, including replications of these experiments on all 10 beers producing a total of 6120 output .mzMLs, which would not be feasible in a lab setting demonstrating the advantage of developing and testing in ViMMS.

## 5 Discussion and Conclusions

Our experiments demonstrated that DDA strategies benefit from RoI-based multi-sample exclusion by collecting equal or more unique fragmentation spectra at higher intensities, and intensity-based exclusion, as expected, improved intensity coverage. Using both, in the single sample case, we saw that we could collect spectra at up to 20% more of the total intensity. The improvements are especially pronounced in the multi-sample case, where we saw that we could collect up to 10% more of the total spectra at up to 20% more of the total available intensity. However, we would expect that when intensity methods reacquire a spectrum at higher intensity they also lose an opportunity to acquire a new spectrum. Although the differences are small, we can observe this in the multi-injection same beer results. But in the mixed-beers 6-4 experiment we see that the intensity methods in fact have slightly better coverage compared to their non-intensity counterpart, and Intensity Non-Overlap has the highest overall.

What causes the coverage increase? Previously in (Davies *et al*., 2021), SmartRoI and Weighted-DEW exchanged some intensity coverage on certain spectra for increased overall coverage against TopN. Although in theory all the scans can be allocated for optimal (intensity) coverage, in a noisy real-world process some degree of redundancy may be desirable. We only see the TopNEXt coverage increase in the multi-sample case, so one potential explanation is that peak-picking has detected different peaks across samples in similar regions of the space, allowing extra coverage when revisiting these locations. A similar argument could be made for SmartRoI variants performing best in our experiments. SmartRoI produces less fragmentation events overall — see the “efficiency” metric in Table 2 of (Davies *et al*., 2021) — so it may create exclusion regions less prematurely in a multi-sample context. This would allow these regions to be visited later compared to other DEW methods, when they would be most relevant. Supplementary Section 5 contains additional experiments which further explore the behaviour of our fragmentation strategies.

Overall, together our simulated and lab results demonstrate that TopNEXt addresses some of the traditional weakness of DDA in terms of sample coverage. By providing more and better quality fragmentation spectra, TopNEXt will aid the process of metabolite annotation, which in turn should lead to greater biological understanding of metabolomics samples. Future work might involve adaptation to different experimental contexts. For example, rather than only being interested in the absolute number of peaks we can acquire spectra for, a specialised method for a case-control setup might value having pairs of spectra from both case and control. Alternatively, we also use RoIs as “peak-like objects” which can be compared for similarity against fragmented objects, but we could instead use a different RoI building algorithm or a more complicated similarity measure than shared area. These developments or others could be flexibly switched out in the TopNEXt framework with the rest of the procedure working as before, easing future fragmentation strategy development. Additionally, the TopNEXt family of methods can be used as-is through ViMMS to perform metabolomics experiments if separate bridging code exists between ViMMS and the mass spectrometry instrument model. This is currently limited to instruments exposing a Thermo Fisher IAPI, but in future bridges to other instrument models may be written.

