## Supplementary information for "TopNEXt: Automatic DDA Exclusion Framework for Multi-Sample Mass Spectrometry Experiments"

#### 1 Beer Data

For our experiments we collected samples of ten arbitrarily-chosen store-bought beers. Store-bought beers are metabolomically complex enough to be a useful test of fragmentation strategy performance, but are also straightforward to obtain and use (alternatives might introduce ethics or bio-safety concerns) which eases reproduction of our results. The list can be found in Table 1. Note that only the first six beers were used for our experiments in the main paper: all ten were used for a simulated replication study which can be found later in this supplementary information.

| Index | Name | Type |
|---|---|---|
| 1 | Raspberry Sour by Vault City | Sour |
| 2 | Cacao & Hazelnut Broken Dream Twisted Breakfast Stout by Siren | Stout |
| 3 | Tennents | Lager |
| 4 | Life and Death by Vocation | IPA |
| 5 | Silence is... Citra by Overtone | Pale Ale |
| 6 | Punk AF by Brewdog | Alcohol-Free IPA |
| 7 | The Hop - Single Hop Series Simcoe Edition by Salt | IPA |
| 8 | 42 DDH Pale Ale by BBNO | Pale Ale |
| 9 | Aoraki by Vocation | DIPA |
| 10 | Punk IPA by Brewdog | IPA |

Table 1: A complete list of the beers collected for our experiments. Note that although the specific choices of sample were arbitrary, we varied the class of beer in the "Type" column in an attempt to obtain varied metabolic profiles.

For the replications of the 6-4 experiment we had each observation randomly sample six beers out of the ten. The random samples we used are in Table 2.

| Observation Num. | Beer Indices |
|---|---|
| 1 | (2, 1, 8, 4, 5, 10) |
| 2 | (9, 4, 5, 7, 10, 1) |
| 3 | (3, 8, 9, 5, 1, 2) |
| 4 | (9, 4, 2, 7, 8, 5) |
| 5 | (5, 6, 4, 7, 9, 2) |
| 6 | (7, 4, 8, 9, 10, 5) |
| 7 | (8, 5, 10, 2, 9, 3) |
| 8 | (7, 4, 5, 10, 1, 2) |
| 9 | (8, 6, 10, 3, 7, 2) |
| 10 | (7, 1, 2, 10, 9, 3) |

Table 2: The random samples used for the 10 replications in the 6-4 replicate experiments.

### 2 Peak-Picking Parameters

#### 2.1 Peak-Picking

To compute our evaluation metrics, we used MZMine 2 (version 2.53) [3] with a fullscan .mzML as the input, producing as output a set of peak-boxes indicating regions we would ideally like to fragment for that sample. For a re-simulated experiment, we used the same fullscan for generation of re-simulations and for creation of peak-boxes. After using the ADAP [2] chromatogram builder and deconvolver and grouping isotopes, we aligned all samples in each experiment. For example, the six different sets of peak-boxes produced by the six different samples in the 6-4 beers experiment. This produces a matrix where rows give a single aligned peak, and columns the sample the peaks were observed in. For each fragmentation run our ViMMS code updates its corresponding column with information like maximum intensities and fragmentation intensities — this allows computing coverage and intensity coverage by collapsing rows and columns.

To produce our steps from chromatogram detection to alignment, we defined an MZMine .xml template file which specifies the order of steps and their parameters (these "batch mode" files can be created and edited through the MZMine GUI). To run this from our codebase we insert the names of the input/output files into a base template .xml, and call MZMine from the command line using Python's subprocess module and this modified template. We have two base templates — "a permissive" parameter set previously used to evaluate SmartRoI and WeightedDEW [1] and a "restrictive" parameter set which was further refined by our mass spectrometry expert. Both of these are available on the ViMMS GitHub repository. The restrictive parameter set filters out lower-quality peaks and more closely represents actual practise, therefore we primarily used this set (including for everything in the main paper) when evaluating our results. The permissive parameter set produces a larger set of peak-boxes and therefore may give us some insight into behaviour on samples more dense with peaks, so we have used it for some additional experiments in this supplementary document. The complete list of batch steps and parameter settings can be found in Tables 3 and 4 — italics are used in the restrictive parameter table to highlight changes in parameter values from the permissive table.

Note that there is one change between our permissive parameter set and the set used for the SmartRoI publication [1]. MZMine requires both a Dalton and ppm value to be specified for m/z tolerances, and will use the maximum of these two values. In order to ensure the ppm value will always be used, we set a dummy value of $10^{(-8)}$ for the absolute tolerance.

| MZMine Parameters (Permissive) | |
|---|---|
| **Raw Data Import** | |
| **Crop Filter** | |
| Retention time | 0.5 to 30 minutes |
| m/z | 50 to 1060 Da |
| **Mass Detection (MS1)** | |
| MS Level | 1 |
| Mass Detector | Centroid |
| Noise Level | 1000 |
| **Mass Detection (MS2)** | |
| MS Level | 2 |
| Mass Detector | Centroid |
| Noise Level | 0 |
| **ADAP Chromatogram Builder** | |
| MS Level | 1 |
| Min group size in # of scans | 3 |
| Group intensity threshold | 500 |
| Min highest intensity | 5000 |
| m/z tolerance | $10^{(-8)}$ Da or 10.0 ppm |
| **Chromatogram Deconvolution** | |
| Algorithm | Wavelets (ADAP) |
| S/N Threshold | 10 |
| S/N Estimator | Intensity window SN |
| Min feature height | 5000 |
| Coefficient/area threshold | 1 |
| Peak duration range | 0.10 to 5.00 |
| rt wavelet range | 0.10 to 1.00 |
| m/z centre calculation | Median |
| m/z range for MS2 scan pairing (Da) | Enabled, 0.01 |
| rt range for MS2 scan pairing (minutes) | Enabled, 0.5 |
| **Isotopic Peaks Grouper** | |
| m/z tolerance | $10^{(-8)}$ Da or 10.0 ppm |
| Retention time tolerance | 0.1, absolute (minutes) |
| Monotonic shape | False |
| Maximum charge | 2 |
| Representative isotope | Most intense |
| **Join Aligner** | |
| m/z tolerance | $10^{(-8)}$ Da or 10.0 ppm |
| Weight for m/z | 80 |
| Retention time tolerance | 0.1, absolute (minutes) |
| Weight for rt | 30 |
| Require same charge state | False |
| Compare isotope pattern | False |
| Compare spectra similarity | False |
| **Export to CSV** | |

Table 3: A table of "permissive" peak-picking parameters used in previous work [1] and for our additional experiments.

| MZMine Parameters (Restrictive) | |
|---|---|
| **Raw Data Import** | |
| **Crop Filter** | |
| Retention time | 0.5 to 30 minutes |
| m/z | 50 to 1060 Da |
| **Mass Detection (MS1)** | |
| MS Level | 1 |
| Mass Detector | Centroid |
| Noise Level | 1000 |
| **Mass Detection (MS2)** | |
| MS Level | 2 |
| Mass Detector | Centroid |
| Noise Level | 0 |
| **ADAP Chromatogram Builder** | |
| MS Level | 1 |
| Min group size in # of scans | 3 |
| *Group intensity threshold* | *2000* |
| Min highest intensity | 5000 |
| *m/z tolerance* | *$10^{(-8)}$ Da or 5.0 ppm* |
| **Chromatogram Deconvolution** | |
| Algorithm | Wavelets (ADAP) |
| *S/N Threshold* | *3* |
| S/N Estimator | Intensity window SN |
| Min feature height | 5000 |
| Coefficient/area threshold | 1 |
| *Peak duration range* | *1.0 to 7.00* |
| *rt wavelet range* | *1.00 to 5.00* |
| m/z centre calculation | Median |
| m/z range for MS2 scan pairing (Da) | Enabled, 0.01 |
| rt range for MS2 scan pairing (minutes) | Enabled, 0.5 |
| **Isotopic Peaks Grouper** | |
| *m/z tolerance* | *$10^{(-8)}$ Da or 5.0 ppm* |
| Retention time tolerance | 0.1, absolute (minutes) |
| Monotonic shape | False |
| Maximum charge | 2 |
| *Representative isotope* | *Lowest m/z* |
| **Join Aligner** | |
| *m/z tolerance* | *$10^{(-8)}$ Da or 5.0 ppm* |
| Weight for m/z | 80 |
| *Retention time tolerance* | *0.5, absolute (minutes)* |
| Weight for rt | 20 |
| Require same charge state | False |
| Compare isotope pattern | False |
| Compare spectra similarity | False |
| **Export to CSV** | |

Table 4: A table of more realistic "restrictive" parameters used for most of our fragmentation strategy evaluations, including those in the main paper.

### 3 Metrics

As we elaborated on in the Methodology section of the main manuscript, *peak coverage* (i.e. coverage) and *intensity coverage* are our two measures of DDA strategy performance. Again, given some set of detected peaks (in our case, from MZMine peak-picking) coverage measures how many detected peaks we have with at least one fragmentation spectra above a minimum intensity threshold. For each peak-box, we award a point to a fragmentation run output for each peak-box which has both a fragmentation event and a valid precursor for that fragmentation event. For each fragmentation event we create a one-dimensional m/z interval (i.e. isolation window) of a user-specified length around its centre, and any peak-box that is completely covered by it on the m/z dimension is considered to be fragmented by it. Any point from the previous MS1 scan above the minimum intensity threshold and within the peak-box is then a precursor to this fragmentation event. The *coverage* is then just the count of peak-boxes for which these two conditions are met. The *cumulative coverage* is the same count with aligned peak-boxes across a multi-injection experiment — each peak is aligned one point if it is fragmented in *any* sample and no more than one for being fragmented across several. For each sample we used the original location of that peak-box in the individual sample, not the aligned summary box. Our results present the (cumulative) coverage as a $[0, 1]$-bounded proportion of the total possible (cumulative) coverage.

*Intensity coverage* generalises this idea by measuring the aggregate quality of these acquisitions. This is done by replacing the binary $0, 1$ score awarded to each peak with a proportion of the maximum fragmentation intensity to the maximum *possible* fragmentation intensity i.e. a score in a $[0, 1]$ range. The maximum fragmentation intensity is just the maximum intensity of any of the fragmentation event's precursors, or 0 if there are none above the minimum threshold; the maximum possible fragmentation intensity is just the maximum intensity of the peak i.e. the maximum intensity of any MS1 points falling within its peak-box in one of the fragmentation runs. We then compute *intensity coverage* and *cumulative intensity coverage* in a similar way to coverage. However, note that when computing maximum intensities for aligned peak-boxes, we use their locations in individual samples but their maximum intensity across all samples.

Finally, note that for all evaluations of a given sequence of fragmentation runs we only include peaks which had a maximum intensity above a minimum threshold in one of those fragmentation runs. This can cause slight inconsistencies between experiment cases (especially if one of them schedules more MS1 scans) but avoids penalising fragmentation strategies for peaks that are completely unobtainable. Furthermore, note that these "window mode" measurements of the coverage and intensity coverage are different from previous work, which used "point mode" [1]. In point mode we treat each fragmentation event as a point rather than an isolation window, and check if the point falls within the peak-box, and assign precursors if they fall within an error tolerance (e.g. 10 ppm).

Coverage and intensity coverage give us a direct measure of how well we are addressing DDA's typical weaknesses. Additionally, we avoid any real-world complications which are not directly relevant to the question we are asking — for example, to meaningfully evaluate any data by metabolite annotation, we would have to have a (relatively) complete and accurate database of metabolites for the sample, whereas evaluating by coverage does not depend on our current knowledge, provided our peak-boxes highlight every region of interest. Even if we do not know which of these peaks correspond to metabolites, if we collect strictly more of them at better qualities, it follows that we will have more correct metabolite identifications, and fewer misidentifications. However, while all our methods presented here build on TopN, so it is unlikely the distribution of peaks they target differs significantly, this remains a potential difficulty. And with intensity coverage, would it be better to collect one peak at 25% of maximum and another at 75%, or to collect two at 50%? Improvements on low-intensity peaks are likely to be more valuable, but intensity coverage will score them equally as long as they are above the fragmentation threshold. Additionally, these metrics cannot make a meaningful comparison to DIA, which can cover all peaks but must then recover usable information from them. Future work will have to compare DDA to pre-scheduling and DIA, but coverage will still be useful in disentangling where differences in metabolite annotations come from.

### 4 Controller Parameters

In order to ensure that the parameters used for dynamic exclusion had reasonable values, we grid-searched a small range of plausible values. These and shared parameter values can be found in the tables below. Highlighted in bold are the values found to be optimal when searching and therefore used for our experiments. The parameters of the actual WeightedDEW implementation slightly differ from the presentation in Section 6.1: rt_tol expresses the total rt-length of window $d_0$ plus $d_1$. We list the values actually used in the code here. Invalid combinations caused by rt_tol having a strictly lesser value than exclusion_t_0 (i.e. $d_0$) were excluded. mz_tol indicates the ppm mass tolerance used for RoI-building/the fixed size of exclusion windows in the case of TopN Exclusion. min_roi_length indicates how many points an RoI must have at minimum before it is discarded.

| **Shared Parameters** | |
|---|---|
| Ionisation mode | Positive |
| Isolation width | 1 |
| Min MS1 intensity ($\lambda_{min}$) | 5000 |
| mz_tol | 10 |
| min_roi_length | 3 |

Table 5: Shared controller parameters that were used throughout all our parameter optimisations and experiments. For the real experiments we instead used an isolation width of 0.7 — this is the minimum our instrument supports.

| **Grid Search Parameter Values** | |
|---|---|
| **Standard DEW** | |
| N | 1, 3, 5, 10, **20** |
| rt_tol | 15, 30, **60**, 120, 240 |
| **SmartRoI** | |
| N | 1, 3, 5, 10, **20** |
| rt_tol | **15** |
| intensity_increase_factor ($\alpha$) | 2, **3**, 5, 10 |
| drop_perc ($\beta$) | 0, $\mathbf{10^{-3}}$, $10^{-2}$, $10^{-1}$ |
| **WeightedDEW** | |
| N | 1, 3, 5, 10, **20** |
| exclusion_t_0 ($d_0$) | **1**, 10, 15, 30, 60 |
| rt_tol ($d_0 + d_1$) | 15, 30, **60**, 120, 240 |

Table 6: Searched parameter values for each DEW variant. All combinations for a given variant were tried: marked in bold are the highest-scoring values, which we used for our actual experiments.

In total there were 227 searched cases. In each case we had ViMMS generate a set of chemical objects from the real fullscan .mzMLs generated for beers 1, 2 and 3. Then for each parameter combination, we ran a smaller version of the 6/4 experiment with 6 total runs following the order 1-2-3-1-2-3. All of these parameter combinations were then sorted by their proportional intensity coverage, and the parameter values used in our actual experiments were the ones with the highest proportional intensity coverage. The fullscans used were generated as part of our lab experiment on the first day, and thereafter we used the parameter values optimised on these. Additional fullscans were generated for use with the fragmentation runs on other days, but the length of the optimisation procedure prevented these being used to optimise parameters for the data generated on the same day.

Although this procedure significantly improved the results across all controllers when comparing re-simulated results (mainly due to increasing the value of the "N" parameter from its initial value of 10) this is not an exhaustive parameter search. As this procedure can be quite time-intensive, the individual domains for the parameters we considered were relatively small, and the 3-2 experiment might behave differently from a 6-4 or 1-20 setup. We found that when ranking these parameter combinations by intensity coverage, WeightedDEW combinations appeared before regular DEW which appeared before SmartRoI. This result was consistent when optimising parameters both on preliminary data and the fullscans generated on day 1 of our lab experiment, so we used WeightedDEW in the lab experiment. However, in our final results,

*SmartRoI* consistently performed the best. We speculate that the different sample setup (3-2 vs 6-4 or 1-20) is the reason for this. Exhaustive comparisons are left to future work.

Additionally, all the controllers share the same dynamic exclusion parameters, so we conducted this search using the Intensity Non-Overlap controller, under the reasoning that it would be best to search using the controller with the most complicated behaviour. However, we could suppose the most dissimilar controller from Intensity Non-Overlap in our comparison, TopN Exclusion, interacts significantly differently with different parameter values. Therefore as a basic sanity check we re-optimised the parameters for the regular DEW using TopN Exclusion instead, and compared TopN Exclusion to Intensity Non-Overlap in a re-simulated experiment. The optimal value of $N = 20$ did not change, but TopN Exclusion instead had a slight preference for $rt\_tol = 30$ rather than a value of 60, so we used this value for both controllers.

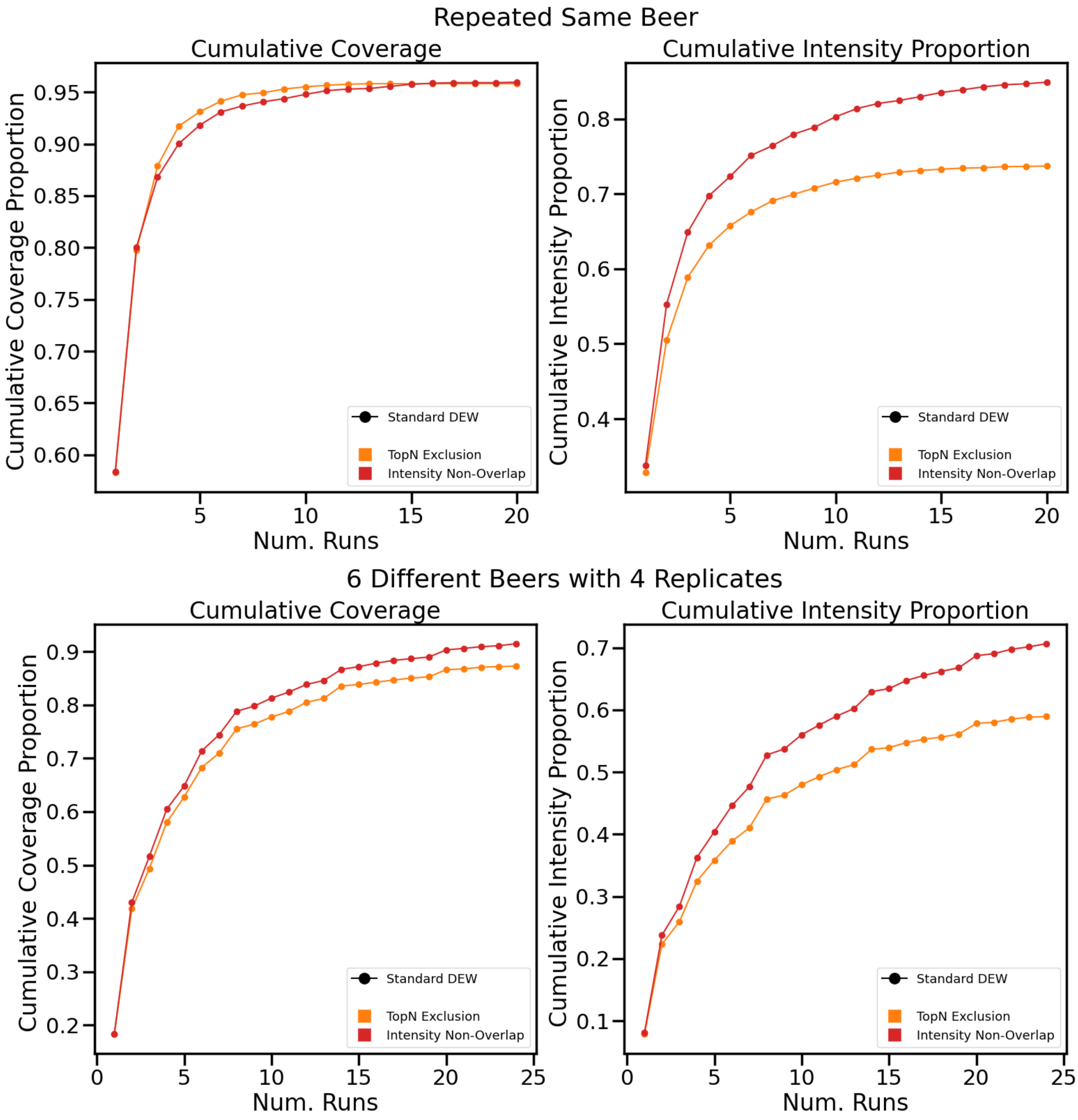


Figure 1: Comparison between TopN Exclusion and regular DEW Intensity Non-Overlap using optimal values for TopN Exclusion and restrictive peak-picking. Top: same beer repeated. Bottom: 6-4 beers.

We can observe in Fig. 1 that this does not significantly change the results of the evaluation. The methods are still competitive in terms of repeated sample coverage, Intensity Non-Overlap has a slight advantage in coverage in the multi-sample case, and Intensity Non-Overlap still has a strong advantage in intensity coverage in both cases.

### 5 Extended Results

All results in the main paper use the same set of real beer data, either from the lab experiment or re-simulated from some of the fullscans produced by it. These were then all evaluated using the restrictive parameter set elaborated on in 2. To give a better idea of the behaviour when generalised to other scenarios, this section contains several alternative experiments.

#### 5.1 Replication Study

It could be argued that the experiments in the main paper may not be reproducible on even our other beer samples, due to only being repeated once. To address this concern we have performed a simulated replication study. Here we repeated both experiments ten times, and aggregated the results per controller — for the same beer experiment we have used a different beer for each replication, and for the 6-4 experiment we have randomly sampled a different set of six beers out of the total set of ten and had each controller perform the 6-4 experiment with these ten sets of six beers. The beers chosen for the 6-4 experiment can be found in Table 2.

In Figures 2 and 3 we see the final coverage and intensity coverage for each of these sets of repeated experiments. The results are highly similar to the individual experiments we have seen in the main paper. For the same beer case, all methods except TopN finished with near-total coverage, and the intensity methods perform best on intensity coverage, with Intensity Non-Overlap having a spike of around 10% compared to non-intensity methods. In the 6-4 case, we again observe that the new methods are a significant improvement on coverage, with Intensity Non-Overlap having a marginal advantage over the others. And for intensity coverage we see again that the new methods performed better, with the Intensity Non-Overlap having the largest advantage. As before, the difference between Intensity Non-Overlap and the TopN Exclusion variants in this case is almost a fifth of the total intensity coverage that can possibly be obtained. In total running these experiments produced $1800 + 4320 = 6120$ .mzMLs across forty-seven hours, using 20 computer cores. For comparison, if we were to run these experiments on our mass spectrometer at 26 minutes per .mzML produced, assuming no downtime between runs, then these experiments would take a total of 110.5 mass spectrometer days. Seeing that these results are consistent under all of these conditions we can therefore be confident that they hold across our selection of beers.

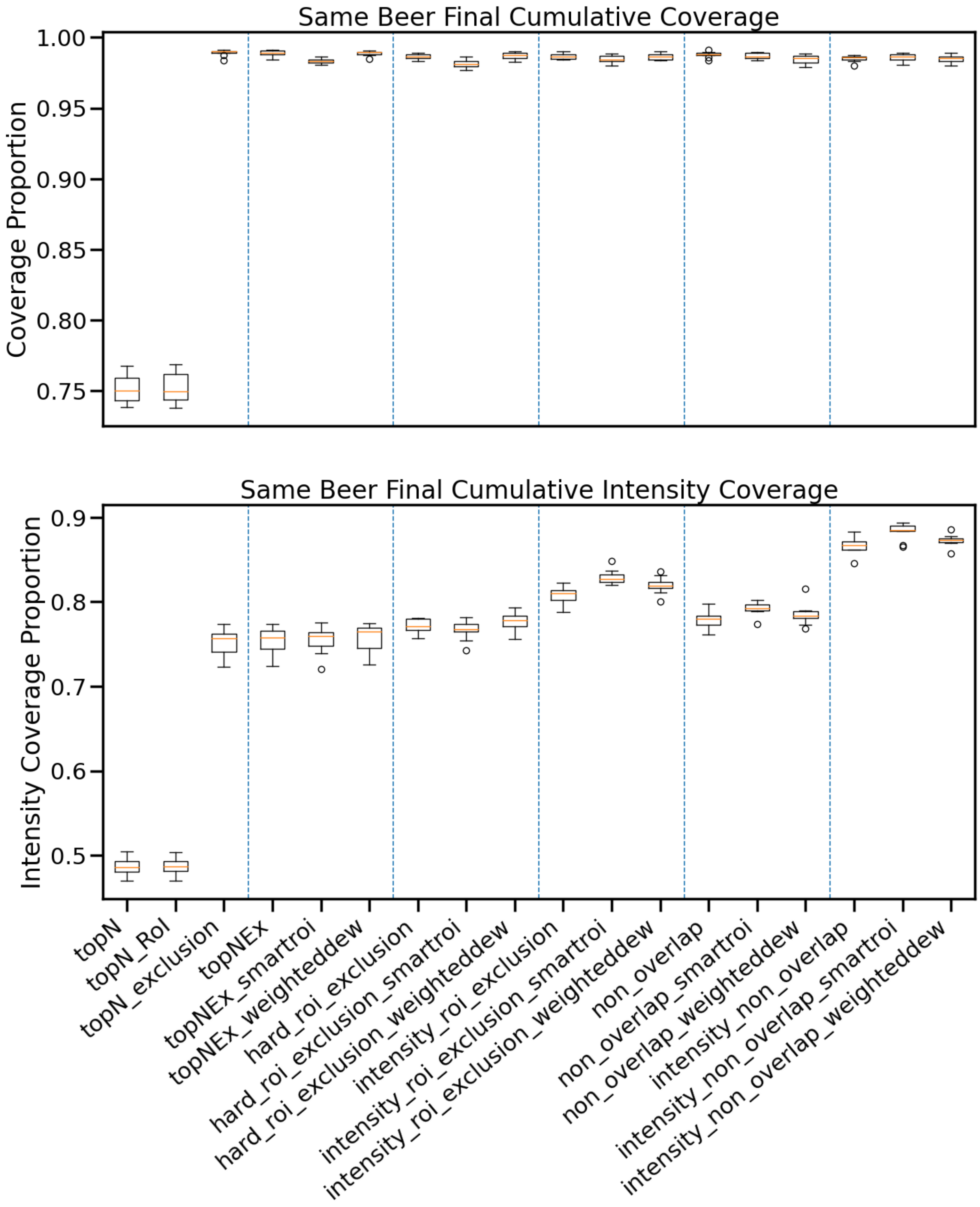


Figure 2: Replicated experiment where for each box there are ten observations, and each observation takes one out of a set of ten different beers (such that all ten beers are each used once per box) and performs a simulated MS experiment with ten repeats of that single beer.

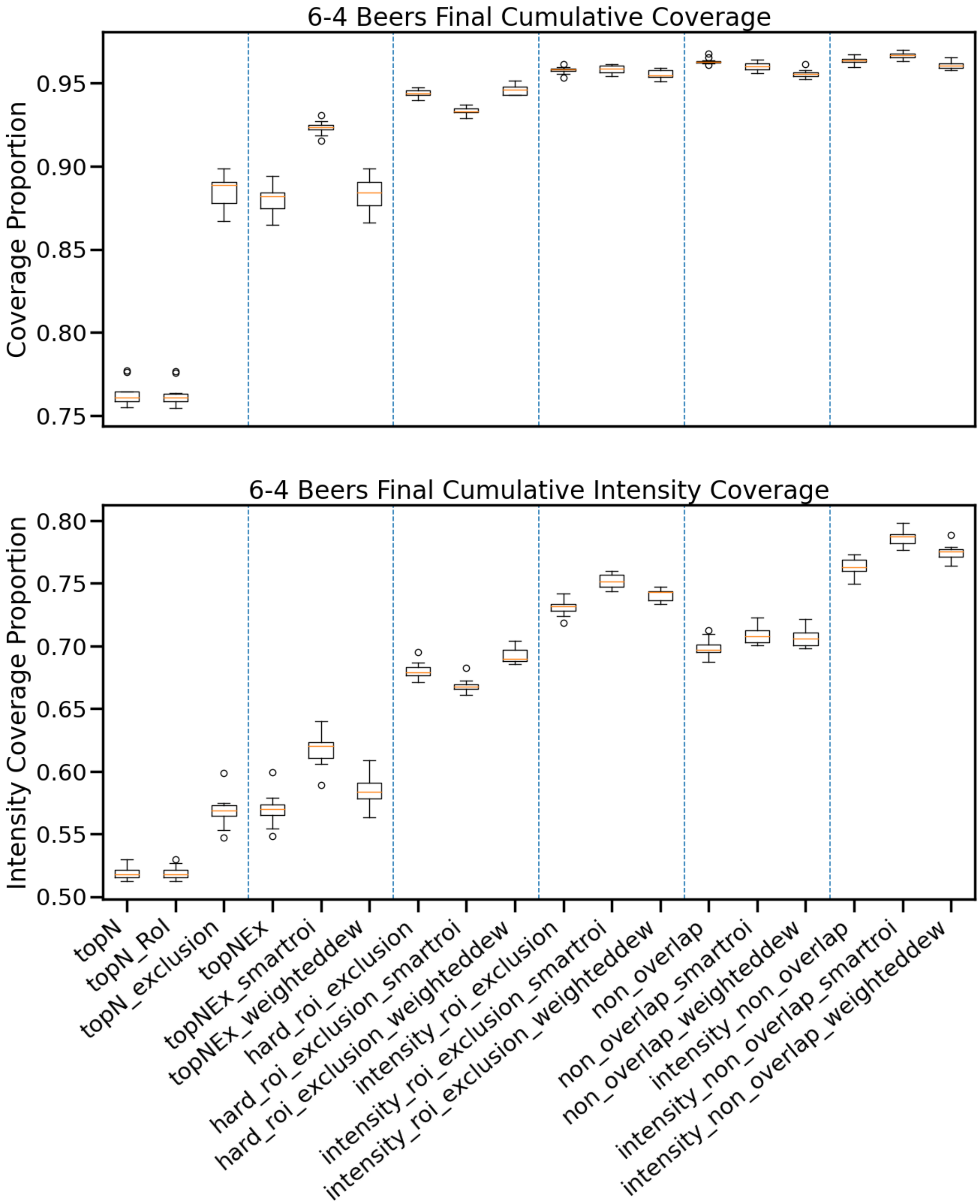


Figure 3: Replicated experiment where for each box there are ten observations and each observation uses a set of six beers that were chosen beforehand by randomly sampling six of the beers, such that each box uses the same set of ten sets of six beers. These six beers are then each repeated four times in a simulated experiment to form the single observation.

#### 5.2 Results with Alternative Peak-Picking

Now we ask what happens if we evaluate the lab results using the permissive peak-picking parameter set described in Section 2 rather than the restrictive set we have used up to this point. The values of the restrictive set were chosen in discussion with our mass spectrometry expert and by design filter out a lot of noise, though they will still likely produce significantly more peaks than there are actual annotatable metabolites. Using the permissive parameter set defines a much greater number of peaks as interesting. This creates a much denser and harder problem, but some of those additional "interesting" peaks may represent a low-intensity metabolite. And, if we could get 100% coverage on the permissive parameter set, we would have done so on the restrictive set as well. Therefore we treat this as a proxy for the behaviour of our controllers on samples that even with a restrictive parameter set still end up dense with interesting peaks.

To give an idea of the actual numbers of peaks involved, restrictive peak-picking on the first beer only produced 2148 identified peaks, but permissive parameters produced 10939. Similarly, for the six beers used in the main 6-4 experiment, restrictive peak-picking parameters produced 6490 identified aligned peaks, and permissive parameters produced 22516.

Figure 4 (top) shows the result of the lab experiment with the same beer repeated 10 times from the main paper, but evaluated using the permissive set of MZMine parameters. This time TopN Exclusion has a noticeable coverage advantage of around 5% over Intensity Non-Overlap, and the intensity coverage disadvantage has shrunk to around 8%. A similar change can be observed with the 6-4 beers evaluated using the permissive parameters in Figure 4 (bottom), where TopN Exclusion has a coverage lead of up to around 3% throughout most of the experiment, but is eventually barely overtaken by the overlap methods near the end of the experiment. Intensity Non-Overlap finishes with only around 7% more intensity coverage. Additionally, we can see that (as we would expect) scores for all methods are lower compared to the restrictive parameter set. For the same beer experiment with the restrictive parameters, all methods quickly obtained comprehensive coverage, with TopN Exclusion effectively no longer increasing in coverage by the 8th sample. With the permissive set, TopN Exclusion (the method with the highest coverage) is only slightly above 80% coverage. With the restrictive set, Intensity Non-Overlap had around 86% intensity coverage, but here has only approximately 63%. A similar result can be seen for the 6-4 beers. TopN has only approximately 49% coverage compared to the approximately 77% it was able to obtain with the restrictive parameters, and has only roughly 35% intensity coverage compared to the restrictive 53%. Similarly the strongest methods only break 70% coverage and 60% intensity coverage compared to the thresholds of 90% and 70% seen with the restrictive parameters.

It is clear from the results in Figure 4 that the permissive parameter set makes the coverage problem significantly harder, and that it gives TopN Exclusion a coverage advantage compared to the new methods. (The performance of the new methods relative to each other has not changed much.) The intensity coverage advantage of the new methods shrinks as well, but it is important to note that intensity coverage is not independent of coverage: if a peak is not covered, then it counts as having an intensity coverage of 0%. In Figure5 we see the intensity coverage of these experiments given that we only include peaks which we have covered for that method in the calculation. That is, this is the average intensity proportion where the denominator uses only peaks we have collected so far rather than the entire dataset of peaks. For the same beer case, we see that the difference in this form of intensity coverage between Intensity Non-Overlap and TopN Exclusion is around 13%, but in the 6-4 case it is only approximately 10%. We can also see that in the 6-4 experiment TopN scores better than anything other than the intensity methods on this alternative intensity coverage measure. This measure simply ignores those peaks which TopN did not cover, which is more than half of the total number. Consequently, the fact that TopN has neglected those peaks to target more fragmentation events at those it did cover benefits TopN on this measure. As we have seen previously [1] having a lot of fragmentation events targeted at a single peak increases the chance that one will fall at the optimal time for fragmentation — whether these fragmentation events target similar locations within-sample or between-samples.

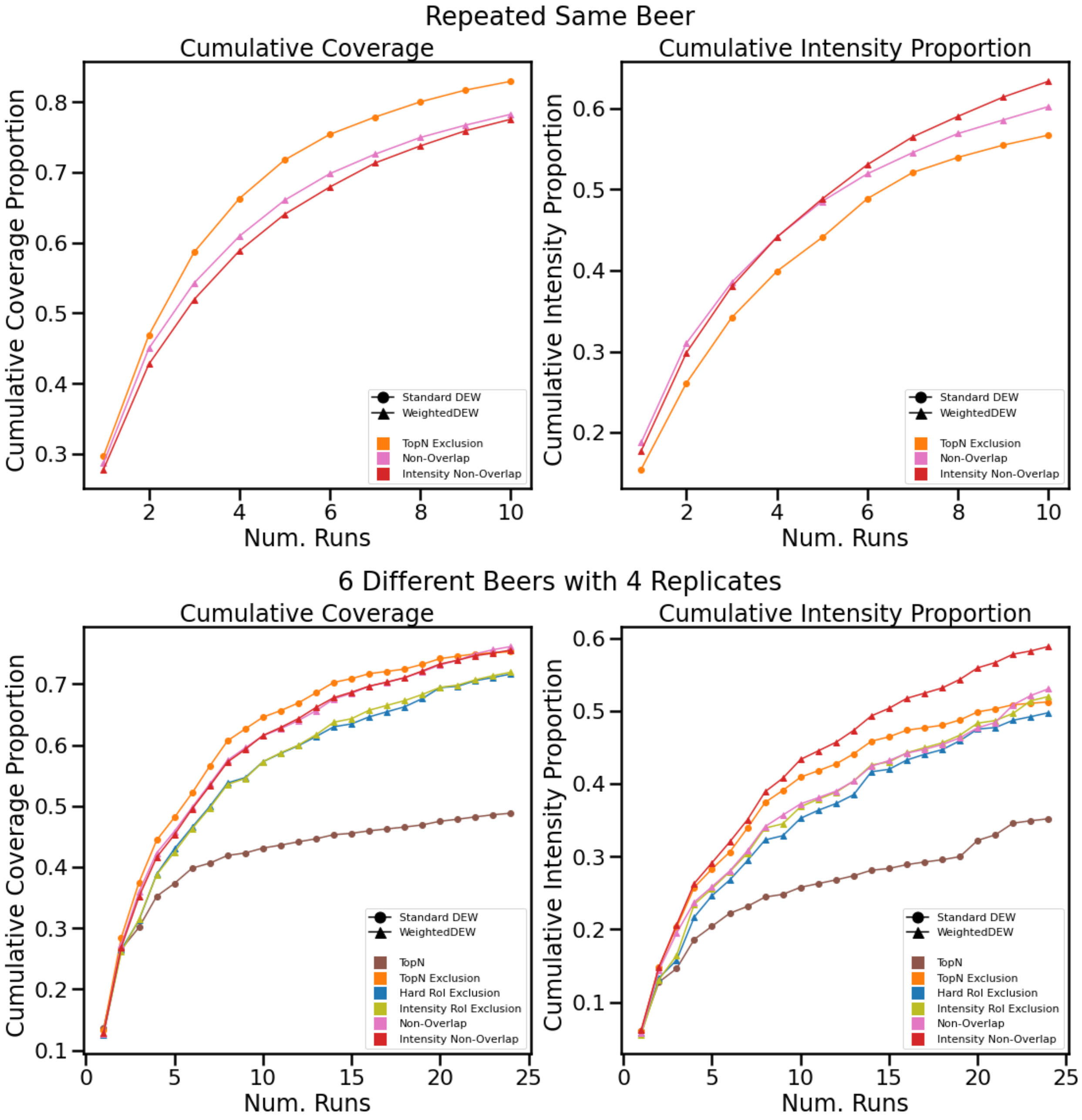


Figure 4: Real experiments evaluated using the permissive parameter set. Top: same beer repeated. Bottom: six different beers each repeated four times.

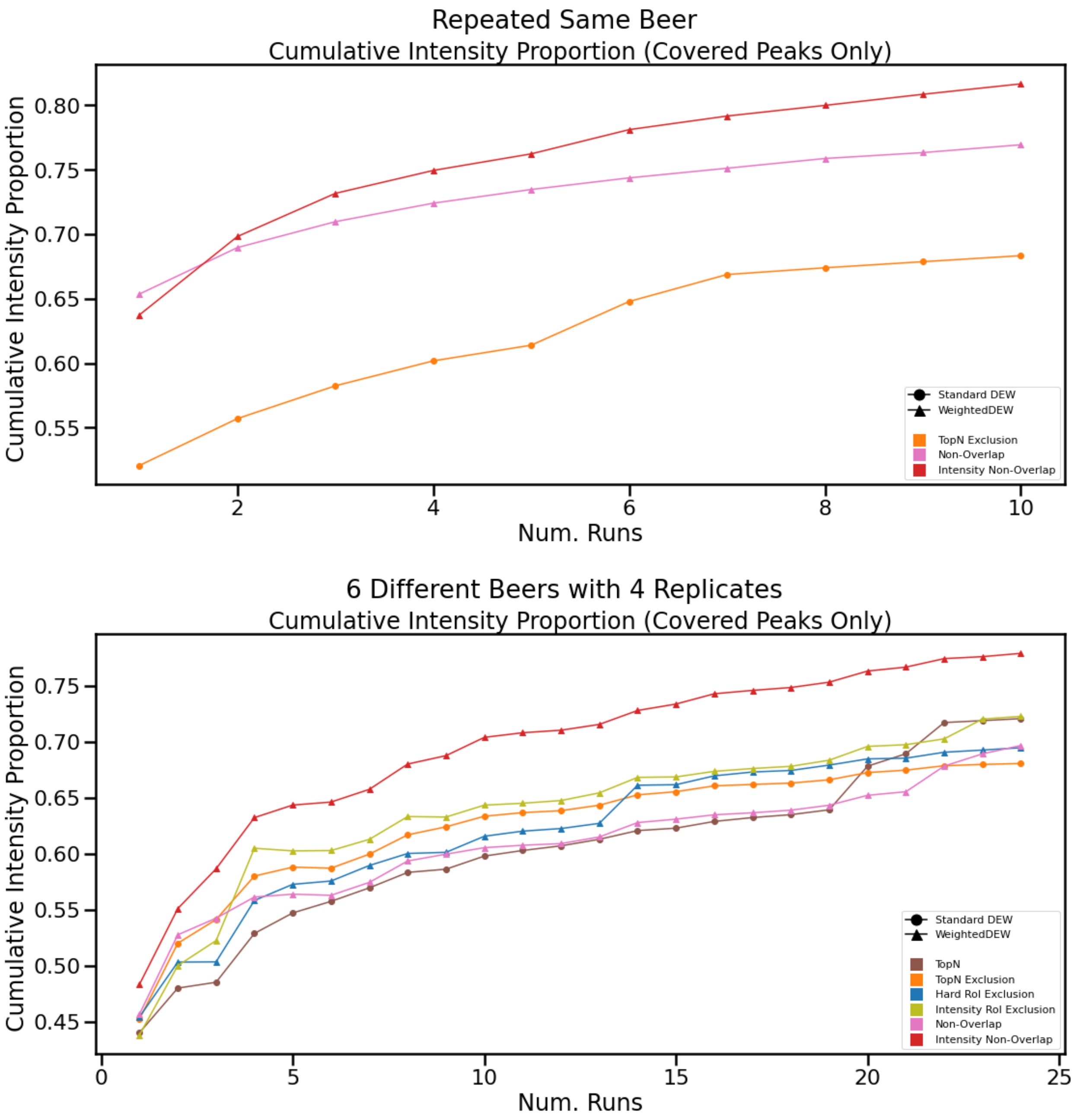


Figure 5: Real experiment evaluated using the permissive parameter set, but showing the intensity proportion where we only include those peaks we have covered. Top: same beer repeated. Bottom: six different beers each repeated four times.

However, even though the relative coverage of Intensity Non-Overlap compared to TopN Exclusion improves from the same beer experiment to the 6-4 beers, the gap in quality of the spectra they do have narrows from 13% to 10%. That is, the spectra we do have with Intensity Non-Overlap are significantly higher quality compared to TopN Exclusion, and this difference is *larger* in the case where the coverage is worse. That most likely indicates that the fragmentation events used to broaden TopN Exclusion's coverage on "weak" peaks not included in the restrictive parameter set are instead being used by the overlap methods to strengthen intensity coverage on the "strong" peaks, even beyond the expected fact that Intensity Non-Overlap attempts to specifically target this behaviour.

Therefore one possible explanation for the results we have seen is that our new methods are more likely to revisit regions of the space compared to TopN Exclusion. By revisiting "strong" peaks instead of visiting "weak" peaks or stopping the duty cycle early and scheduling another MS1 scan, we increase the chance that get a higher intensity coverage on those "strong" peaks or obtain similar peaks across multiple samples. If relatively "strong" peaks are the only things included in the evaluation (as with the restrictive parameter set) then naturally this confers an advantage.

To further substantiate the hypothesis that the new methods revisit potential peaks more often, Tables 7 and 8 show counts of fragmentation events per controller. As the number of runs increases, the number of valid targets for fragmentation decreases and so MS1 scans are scheduled instead of MS2 scans. In the simulated experiments we can see that over the course of the experiment TopN Exclusion eventually almost entirely stops scheduling new fragmentation events, while Non-Overlap decreases significantly and Intensity Non-Overlap remains nearly constant. The likely implication of this is that TopN Exclusion is running out of targets because it does not revisit them, whereas Non-Overlap and Intensity Non-Overlap do and thus continue to have targets to schedule fragmentation events on. One further piece of evidence is shown in Figures 6 and7 which show counts of the number of injections each peak was fragmented in for both experiments with both peak-picking parameter sets. In each case it can be seen that fragmentation events are distributed more heavily towards repeated fragmentations for Non-Overlap and especially Intensity Non-Overlap. This confirms that these methods are more likely to revisit previously fragmented peaks, and together with the fragmentation counts suggests that TopN Exclusion neglects doing this in favour of scheduling MS1 scans.

Another possible explanation is that because TopN Exclusion more often schedules MS1 scans instead of MS2, then it is more likely to see opportunities where "borderline" peaks are the most appealing option. Also, while the evaluation for an individual controller does not include any peaks that do not have a single eligible precursor above the intensity threshold in one of the fragmentation runs tied to that controller, it is also still possible that the permissive parameter set allows many artifacts or otherwise unreachable peaks. Additional MS1 scans may allow these to be considered for fragmentation by the fragmentation strategy.

Based on these results we might tentatively conclude that the new methods are much more thorough in exploring relatively "strong" peaks compared to TopN Exclusion, and thus will get higher intensity coverage on these in particular, as well as higher intensity coverage in general. In multi-sample cases (rather than just multi-injection cases with the same sample) revisiting may also confer an advantage in coverage. However, in cases where we must very rapidly obtain sample coverage on a very dense sample without caring so much about the intensity coverage, TopN Exclusion may be preferable. Trying to find the right balance of broadening coverage vs revisiting in hopes of increasing coverage/intensity coverage may guide the development of future controllers. However, while this evidence may give us some indication of the final behaviour of our controllers, it is insufficient by itself. A real example of a sample dense with interesting peaks likely has a very different distribution of intensities. Therefore in this case we might also expect Intensity Non-Overlap, etc to revisit the same targets less frequently, also broadening their coverage. Further work is needed to determine the specifics of the behaviour in this case.

| Run Index | TopN Exclusion | Non-Overlap | Intensity Non-Overlap |
|---|---|---|---|
| Same Beer | | | |
| 1 | 6293 | 6294 | 6433 |
| 2 | 3556 | 5846 | 6485 |
| 3 | 2590 | 5382 | 6384 |
| 4 | 2186 | 5101 | 6320 |
| 5 | 1839 | 5030 | 6289 |
| 6 | 1589 | 4692 | 6331 |
| 7 | 1388 | 4530 | 6306 |
| 8 | 1126 | 4519 | 6299 |
| 9 | 879 | 4228 | 6254 |
| 10 | 619 | 4140 | 6269 |
| 11 | 467 | 4227 | 6268 |
| 12 | 308 | 4118 | 6229 |
| 13 | 218 | 4048 | 6159 |
| 14 | 198 | 3694 | 6226 |
| 15 | 147 | 3659 | 6188 |
| 16 | 123 | 3791 | 6186 |
| 17 | 108 | 3460 | 6257 |
| 18 | 95 | 3396 | 6235 |
| 19 | 81 | 3296 | 6255 |
| 20 | 79 | 3299 | 6173 |
| 6-4 Beers | | | |
| 1 | 6293 | 6294 | 6294 |
| 2 | 4470 | 5966 | 6178 |
| 3 | 3193 | 5817 | 6185 |
| 4 | 3256 | 5274 | 6031 |
| 5 | 2397 | 5118 | 6064 |
| 6 | 2227 | 4973 | 5942 |
| 7 | 2244 | 4758 | 5970 |
| 8 | 2464 | 4756 | 6104 |
| 9 | 1447 | 4570 | 6093 |
| 10 | 2021 | 4508 | 6025 |
| 11 | 1431 | 4301 | 5947 |
| 12 | 806 | 4236 | 5863 |
| 13 | 1310 | 4036 | 5875 |
| 14 | 1789 | 4240 | 6004 |
| 15 | 518 | 4041 | 5982 |
| 16 | 1287 | 4077 | 6015 |
| 17 | 752 | 3809 | 5969 |
| 18 | 221 | 3763 | 5791 |
| 19 | 686 | 3729 | 5866 |
| 20 | 1306 | 4099 | 6071 |
| 21 | 177 | 3701 | 5921 |
| 22 | 748 | 3827 | 5898 |
| 23 | 333 | 3570 | 5948 |
| 24 | 104 | 3657 | 5744 |

Table 7: Number of fragmentation events per controller during the simulated experiment. Note that Non-Overlap and Intensity Non-Overlap are using the WeightedDEW variant for comparison to the lab experiment.

| **Run Index** | **TopN Exclusion** | **Non-Overlap** | **Intensity Non-Overlap** |
|---|---|---|---|
| Same Beer | | | |
| 1 | 6330 | 5941 | 5922 |
| 2 | 4124 | 5863 | 5918 |
| 3 | 3199 | 5848 | 5918 |
| 4 | 2823 | 5866 | 5857 |
| 5 | 2559 | 5858 | 5891 |
| 6 | 4262 | 5833 | 5901 |
| 7 | 3347 | 5812 | 5821 |
| 8 | 2277 | 5641 | 5861 |
| 9 | 2253 | 5864 | 5839 |
| 10 | 2198 | 5692 | 5808 |
| 6-4 Beers | | | |
| 1 | 6350 | 5909 | 5819 |
| 2 | 5038 | 5914 | 5903 |
| 3 | 3568 | 5918 | 5915 |
| 4 | 3374 | 5923 | 5959 |
| 5 | 2804 | 5945 | 5924 |
| 6 | 2504 | 5843 | 5823 |
| 7 | 2744 | 5881 | 5850 |
| 8 | 2993 | 5807 | 5836 |
| 9 | 2508 | 5851 | 5849 |
| 10 | 2609 | 5874 | 5880 |
| 11 | 2318 | 5892 | 5874 |
| 12 | 1856 | 5776 | 5718 |
| 13 | 2307 | 5834 | 5776 |
| 14 | 2702 | 5851 | 5761 |
| 15 | 2272 | 5861 | 5783 |
| 16 | 2488 | 5888 | 5811 |
| 17 | 2160 | 5776 | 5778 |
| 18 | 1245 | 5793 | 5632 |
| 19 | 2169 | 5767 | 5681 |
| 20 | 2553 | 5800 | 5690 |
| 21 | 2144 | 5737 | 5673 |
| 22 | 2326 | 6188 | 5728 |
| 23 | 1960 | 6183 | 5709 |
| 24 | 960 | 6091 | 5534 |

Table 8: Number of fragmentation events per controller during the lab experiment.

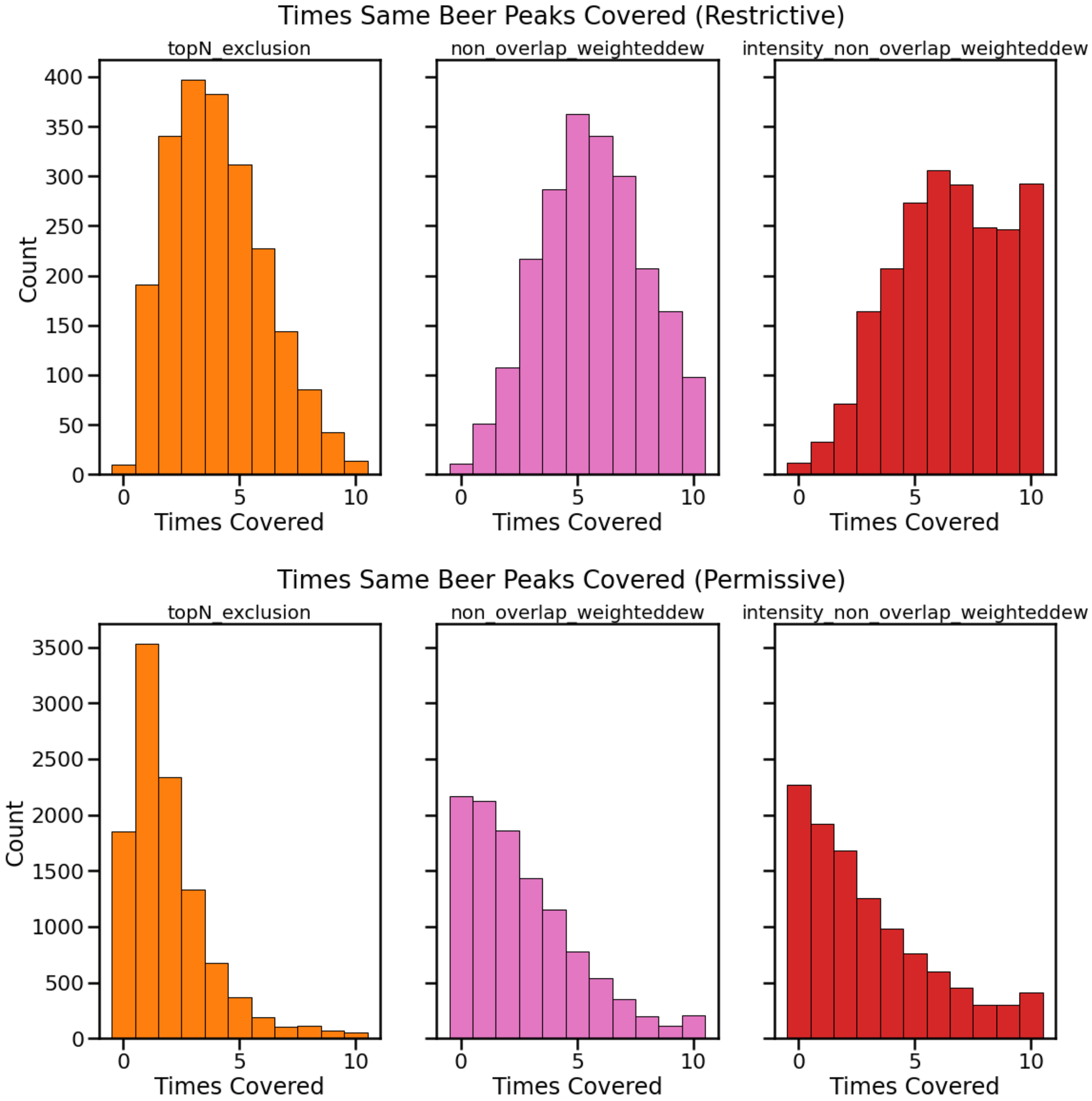


Figure 6: Counts of number of different injections each peak was fragmented in where the same beer was repeated ten times. Top: restrictive peak-picking. Bottom: permissive peak-picking.

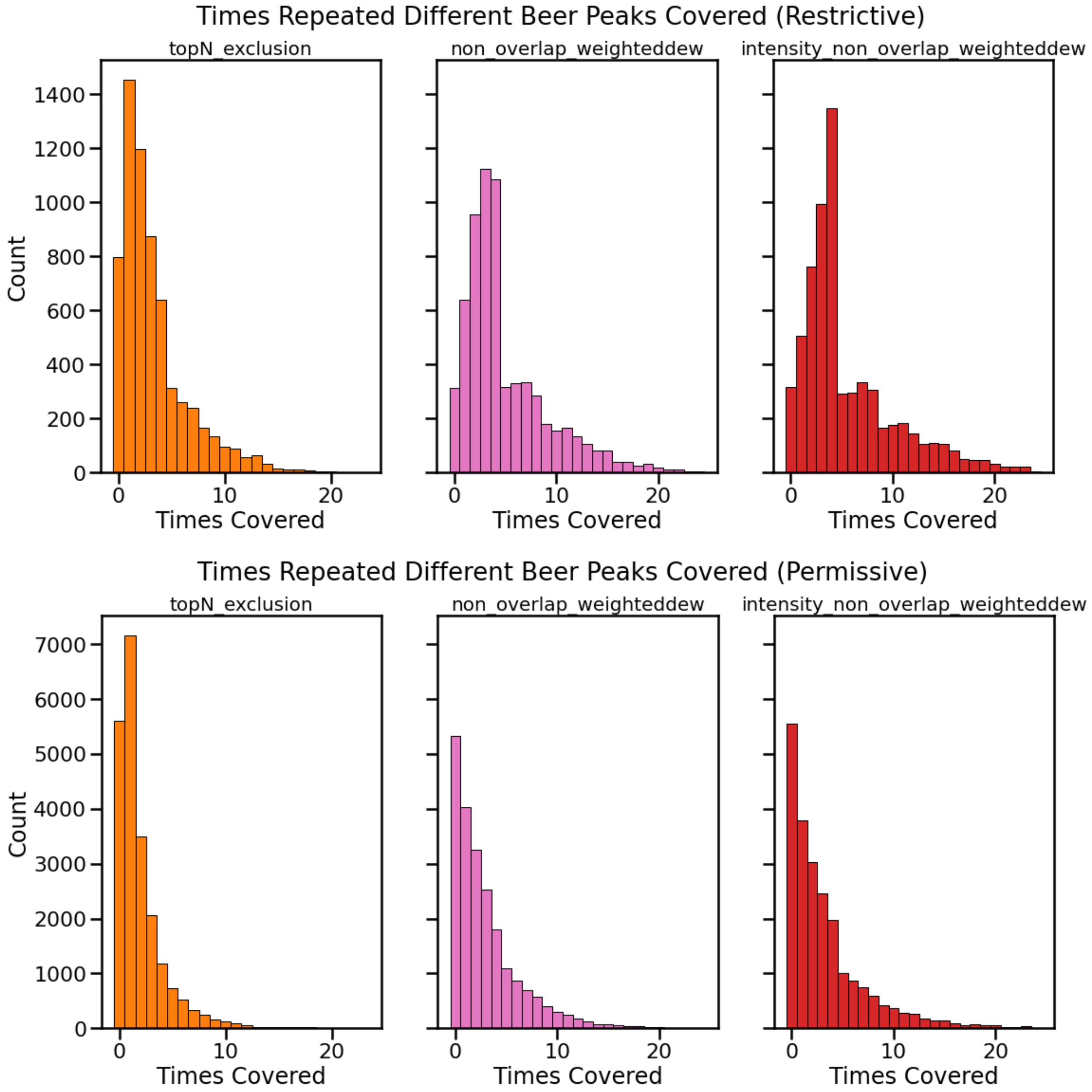


Figure 7: Counts of number of different injections each peak was fragmented in where four beers were each repeated six times. Top: restrictive peak-picking. Bottom: permissive peak-picking.

#### 5.3 Extra Beers+Urines

Having drawn all results to this point from the same pool of data, we now ask about how performance generalises on different data. While we cannot feasibly run another large-scale lab experiment, we can re-simulate any data we have fullscans for. For this we use a previously published data set [4], which we used to test our methods prior to having any lab results to generate re-simulated data from. Although this dataset also uses beers, they are a different set from those in Table 1, and were run in a significantly different experimental context, with a different mass spectrometer (Q-Exactive) and with different instrument settings — the details can be found in the original publication [4]. We again fixed the scan lengths of our simulations to the average scan lengths of the real TopN runs, 0.28 (MS1) and 0.13 (MS2).

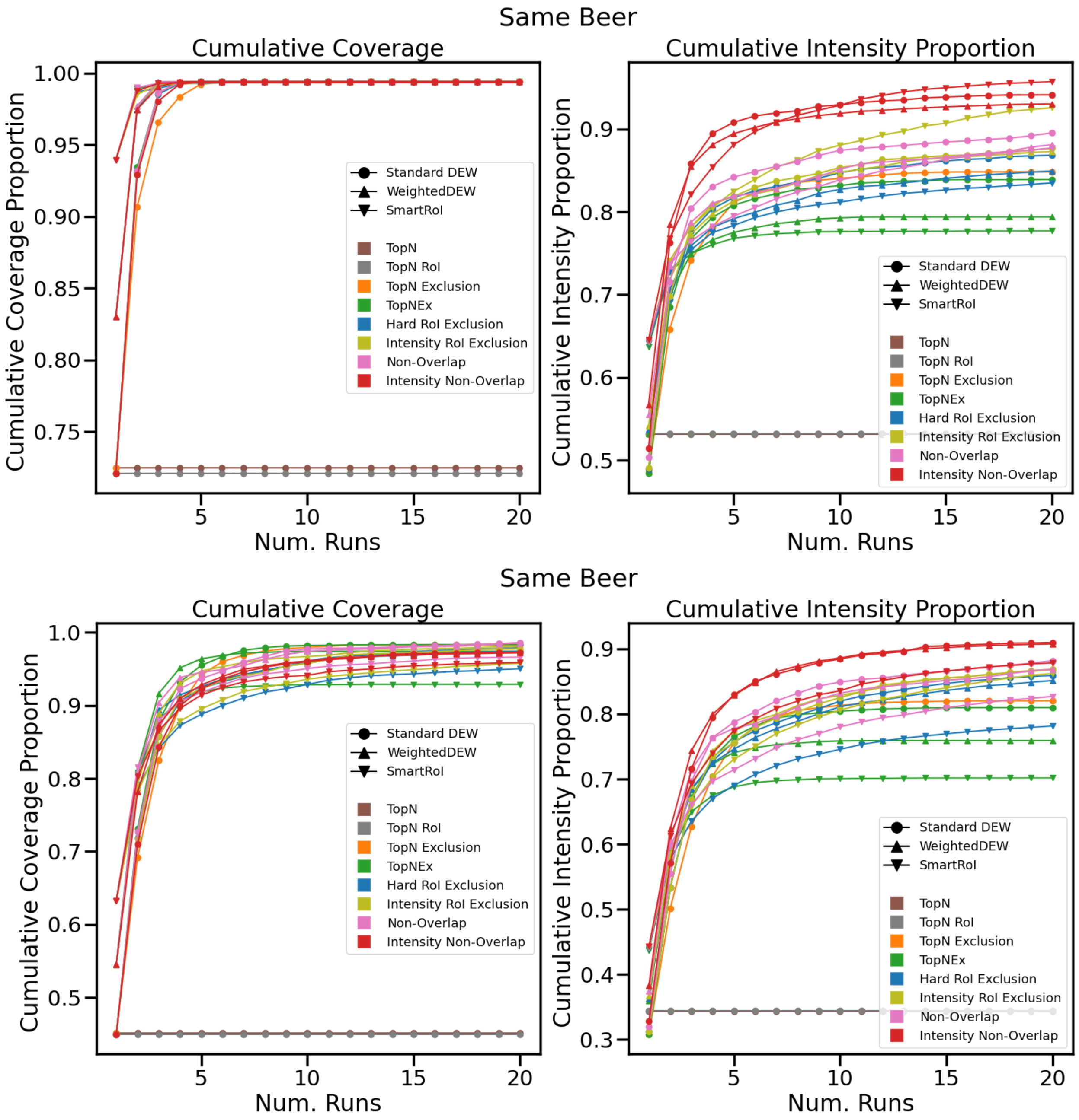


Figure 8: Simulated experiment using the alternative beers with the same beer repeated for ten injections. Top: restrictive peak-picking. Bottom: permissive peak-picking.

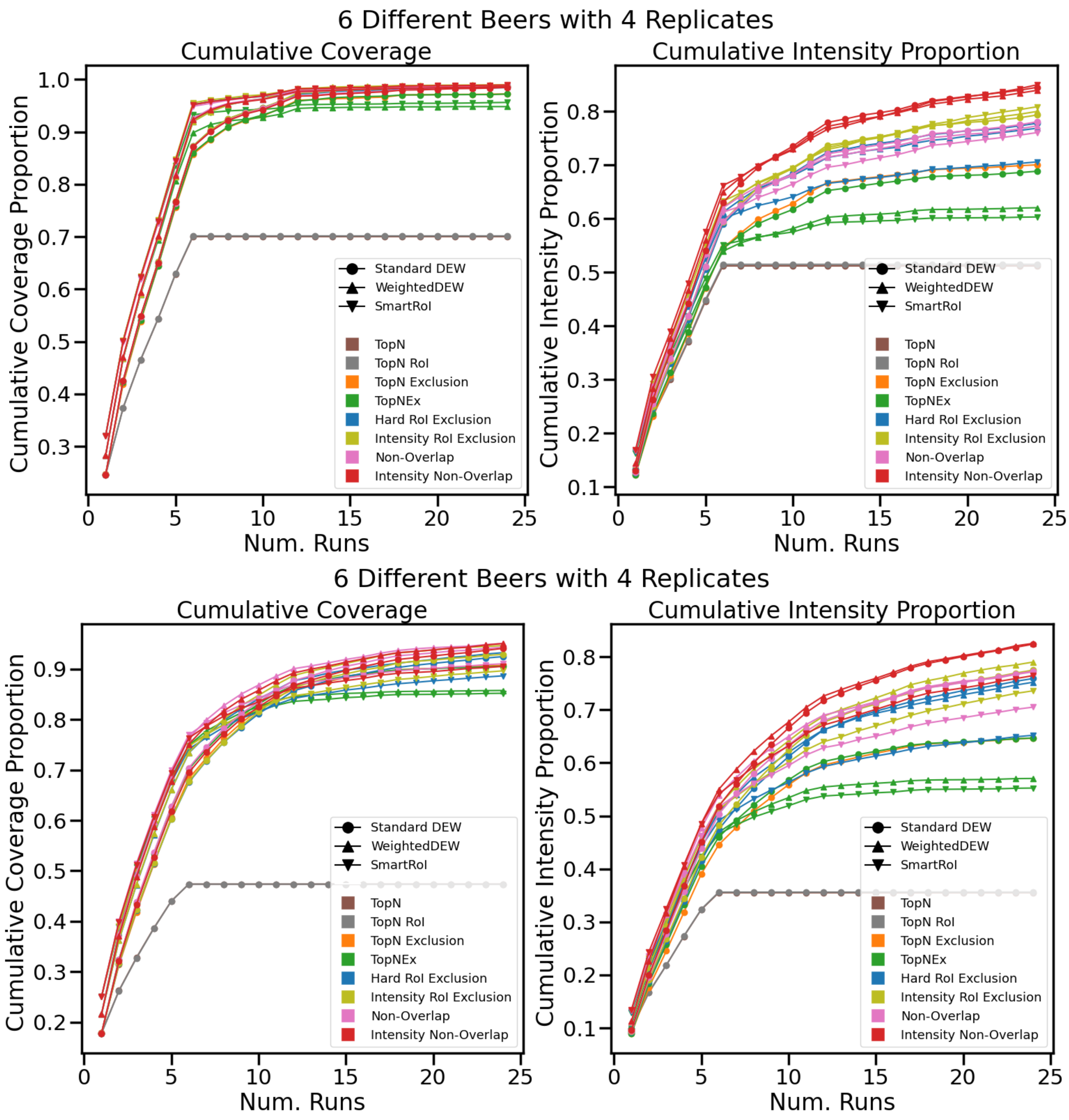


Figure 9: Simulated experiment using the alternative beers with six different beers each repeated four times. Top: restrictive peak-picking. Bottom: permissive peak-picking.

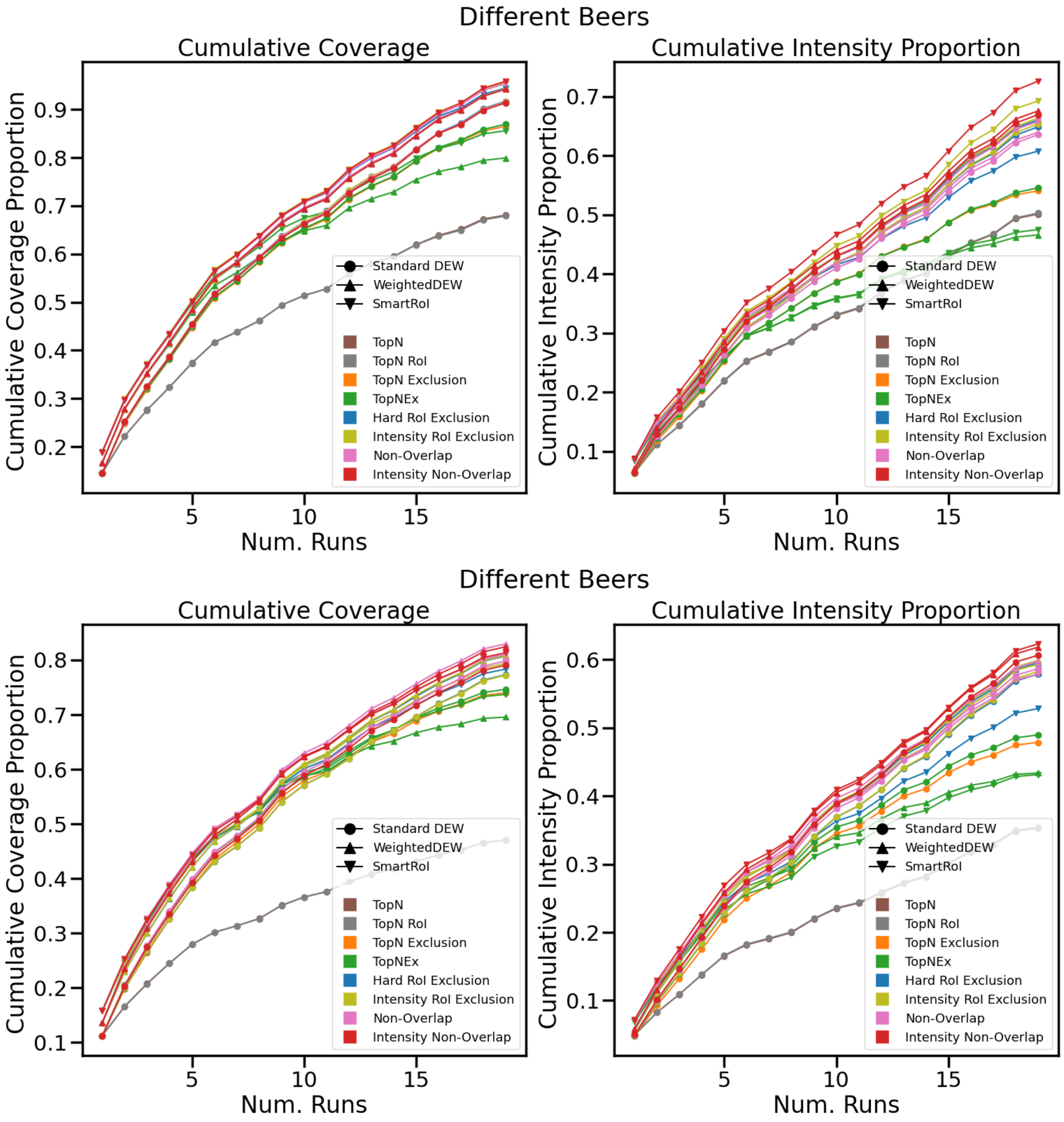


Figure 10: Simulated experiment using the alternative beers with nineteen different beers run once each. Top: restrictive peak-picking. Bottom: permissive peak-picking.

Figures 8 and 9 show results on these beers for both the same beers and 6-4 beers cases we have already explored. Figure 10 leverages the fact that this dataset contains 19 beers to perform a new kind of experiment where each beer is only run once. The equivalent with only six beers would be to read the 6-4 plot up to the sixth injection. The top plots in Figures 8, 9 and 10 show the results evaluated with the restrictive peak-picking parameters; the bottom plots show them with the permissive parameters. The same main conclusions can be observed once again. TopN only gains performance while new samples are injected, and overall performs significantly worse than multi-sample methods even prior to this point. Intensity methods easily perform best in intensity coverage, with Intensity Non-Overlap clearly dominant in all six cases. One notable difference is that although we observed in Section 5.2 that the permissive parameter set increased the coverage of the TopN Exclusion variants relative to the new methods, the same pattern is not as pronounced here. In fact, it seems to only be clearly exhibited in the same beer case, but even in this case the Non-Overlap methods are close in performance, and in the other cases they are significantly stronger. It would therefore seem that this effect generated by the permissive parameters and the overall performance of the controllers are significantly dependent on the input data, and perhaps future work can isolate the reason why, to guide appropriate method use. Peak-peaking for the same beer case produced 1705 identified peaks with the restrictive parameters with the restrictive parameters, and 6901 with the permissive. For the 6-4 case, 5004 restrictive and 17471 permissive identified aligned peaks were produced. For the 19 different beers, 8474 restrictive and 27730 permissive identified aligned peaks were produced.

However, this dataset does not *only* contain beers. It also contains a number of human urines run in the same experimental context — now we can investigate the performance of our methods on a different kind of metabolic sample. Figures 11, 12 and 13 show the same experiments, but with urines instead of beers, and using 15 urines in the different urine case compared to 19 beers. Again, the top plots in Figures 11, 12 and 13 show the results evaluated with the restrictive peak-picking parameters; the bottom plots show them with the permissive parameters. Despite using a different type of sample these results have a very similar profile to the beer results we have just seen, and the same conclusions seem to apply, though notably the best DEW variant seems to vary more strongly between them. Overall this suggests that these results may generalise to other kinds of data and experimental setups, and that the advantages of Intensity Non-Overlap are consistent.

Peak-peaking for the same urine case produced 1142 identified peaks with the restrictive parameters with the restrictive parameters, and 5531 with the permissive. For the 6-4 case, 3903 restrictive and 15264 permissive identified aligned peaks were produced. For the 15 different urines, 7389 restrictive and 26682 permissive identified aligned peaks were produced.

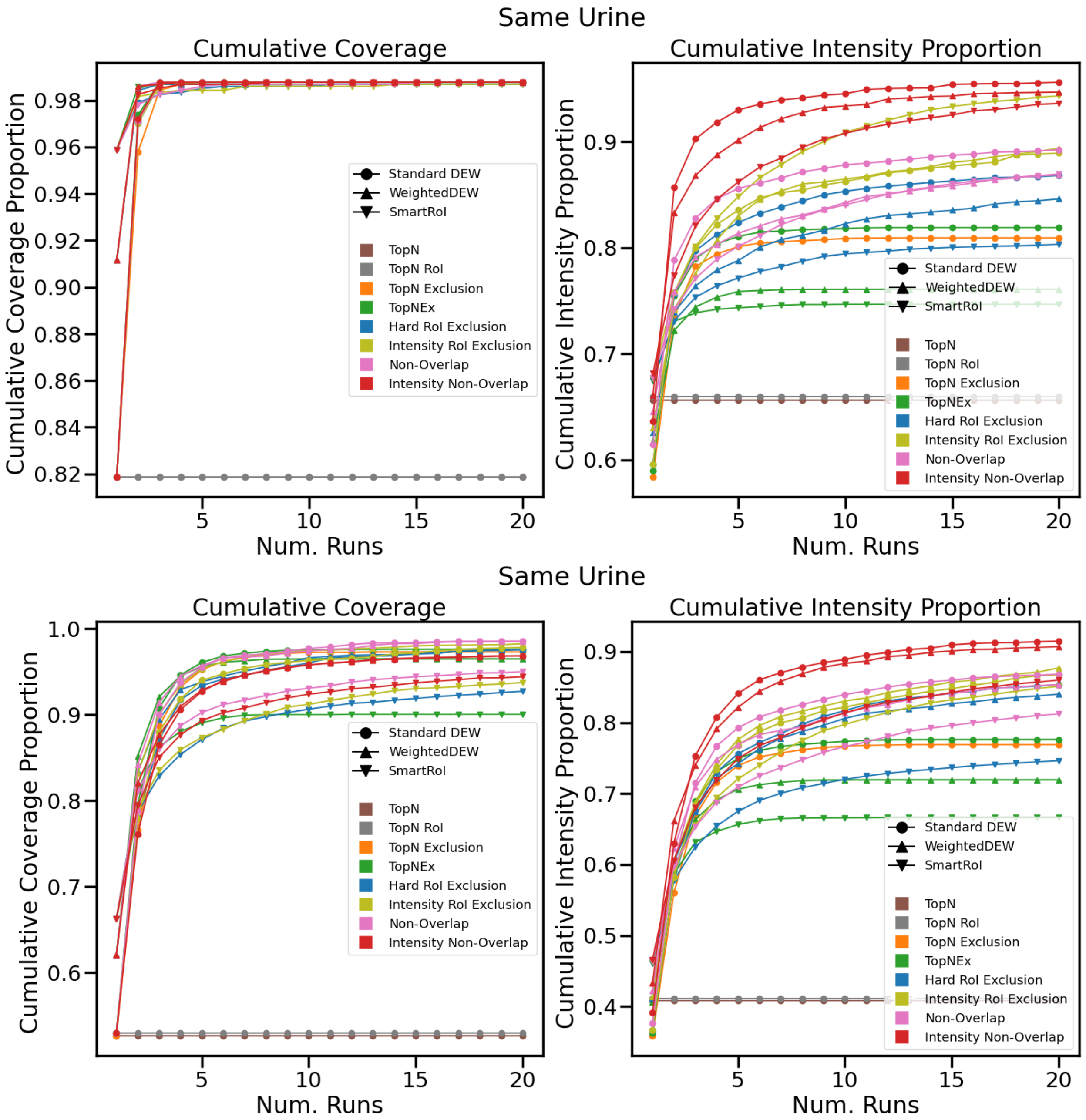


Figure 11: Simulated experiment with the same urine repeated for ten injections. Top: restrictive peak-picking. Bottom: permissive peak-picking.

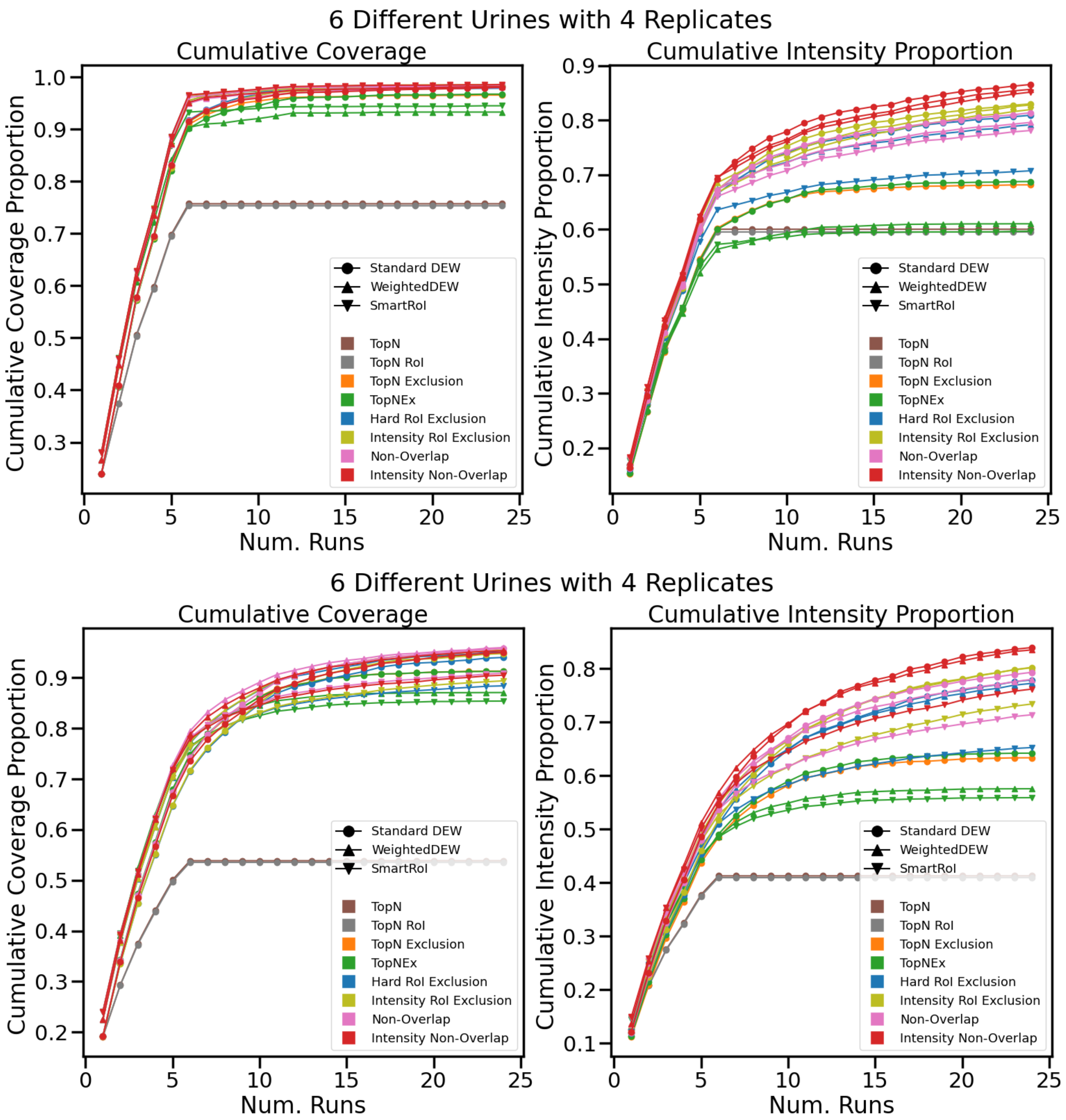


Figure 12: Simulated experiment with six different urines each repeated four times. Top: restrictive peak-picking. Bottom: permissive peak-picking.

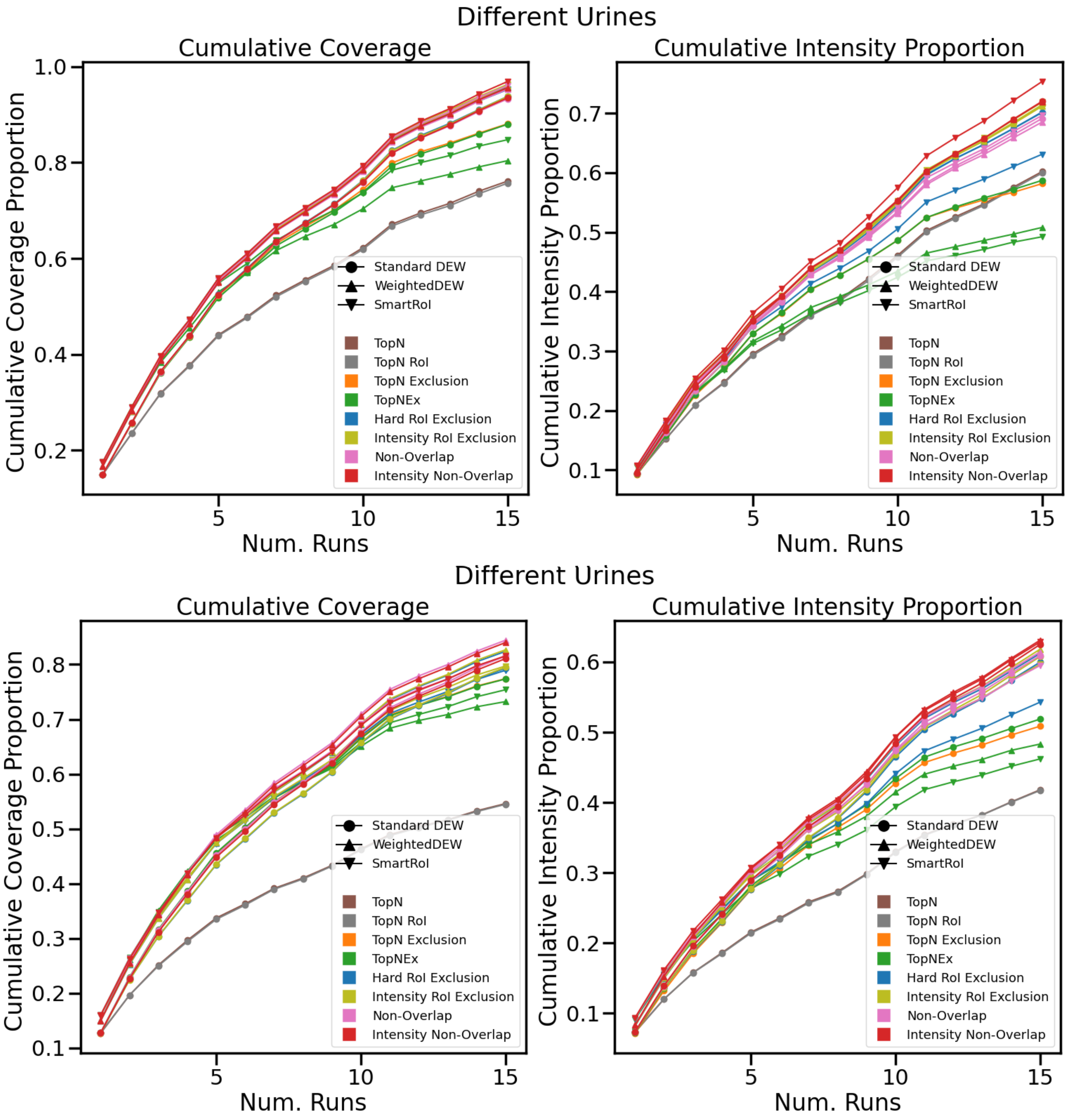


Figure 13: Simulated experiment with fifteen different urines run once each. Top: restrictive peak-picking. Bottom: permissive peak-picking.

### 6 Algorithms

#### 6.1 DEW Indicators

In WeightedDEW, a DEW is extended to be split into two adjacent time intervals with the first beginning at $t_f$ and with lengths $d_0$ and $d_1$. In the first interval, normal DEW exclusion occurs. In the second, a $[0, 1]$-bounded weight is instead applied to the intensity, equal to the proportion of the rt-distance the precursor has from the second interval's starting-point $t_f + d_0$. Equation 1 shows $f_{ex}(p, Ex)$, which can be directly substituted for $I_{ex}$ — note that our implementation applies this formula only to DEWs and not to multi-sample exclusion windows, which act as before.

$$f_{ex}(p, Ex) = \begin{cases} 1 & \text{if } p \text{ is not within a DEW} \\ \frac{t_p - (t_f + d_0)}{d_1} & (t_f + d_0) < t_p < (t_f + d_0 + d_1) \\ 0 & t_f < t_p \leq (t_f + d_0) \end{cases} \tag{1}$$

We found that WeightedDEW weights were sensitive to changes in their relationship with logarithms in e.g. Intensity RoI Exclusion (similarly to non-overlap, the weight worked best as a power on the unlogged modified intensity) so this motivated the placements of many of the logarithms in our equations.

SmartRoI builds on RoI-based DEWs by replacing the regular DEW with a system built on two new parameters, $\alpha$ and $\beta$. $\alpha$ controls the proportion by which a RoI's intensity must be higher than the previous fragmentation intensity $\lambda_f$: $\beta$ controls the proportion of $\lambda_{max}$ it must have fallen below, where $\lambda_{max}$ is the highest point since the last fragmentation. If $\lambda_r$ has not changed by more than either of these thresholds, $r$ is excluded from fragmentation. Equation 2 shows $I_s(r)$ which can be substituted in the same way as $f_{ex}(p, Ex)$.

$$I_s(r) = \begin{cases} 0 & \text{if } r \text{ has been fragmented and } (\frac{\lambda_r}{\lambda_f} < \alpha) \text{ and } (\frac{\lambda_r}{\lambda_{max}} > \beta) \\ 1 & \text{otherwise} \end{cases} \tag{2}$$

#### 6.2 Algorithmic Details

Our controllers for our new fragmentation strategies rely on some simple computational geometry. In all cases, for each active RoI in the current scan, we must find all non-DEW exclusion windows related to it, so that we can then compute a scoring function for it. TopN Exclusion, Hard RoI Exclusion and Intensity RoI Exclusion all treat the last precursor in the RoI as a point, then ask which exclusion windows it falls into. TopN Exclusion and Hard RoI Exclusion only want the boolean value of whether the precursor falls into any window, to compute the indicator function: Intensity RoI Exclusion must know all of the windows the precursor falls into so that it can retrieve the maximum of their intensities. For Non-Overlap and Intensity Non-Overlap, we must retrieve all exclusion windows overlapping a given query RoI to extract their shared area.

##### 6.2.1 Filtering Exclusion Windows

For all controllers we must perform this search in real-time for every precursor in a given MS1 scan. An MS1 scan takes around 0.6 seconds on our instrument, so we would ideally like processing time to be on the order of $10^{(-2)}$ or less. However, given that over the course of 20 successive injections the number of exclusion windows can reach the 100,000s, and we must search this set of windows for *every* precursor in the current scan, a naive containment check where we simply iterate through every (precursor, window) pair is not practical. Therefore we separate the space into a discrete grid of fixed-size boxes, and each of these grid-box stores which of the exclusion windows overlap it. Simple arithmetic then computes which of these grid-boxes a precursor falls into, and then we need only manually check that the precursor is contained within the small subset of exclusion windows associated with its grid-box. This data structure is updated entirely offline, between injections, when all exclusion windows for that injection are added to the total.

A simple example is to imagine dividing the space into four, then for each precursor only checking the exclusion windows overlapping the quadrant the precursor falls into. If the exclusion windows were evenly distributed, then this would divide the number needing to be searched by four. However, when querying an entire RoI for area exclusion it may lie across multiple grid-boxes, and we must take the union of their contents before checking them for intersection. But in our simple example, even if the query RoI lay across two quadrants, then we would still only need to search half the number of exclusion windows for intersection.

By quartering the space, we have split it once on each dimension, but we extend this idea and split it more times to reduce the set of containment/intersection checks further. However, more splits is not always better. Increasing the number of grid-boxes also creates a memory overhead (and processing overhead in managing that memory) and making the grid-boxes smaller also increases the number of them that exclusion windows will be duplicated between, requiring additional processing for the query RoI to filter them down into a single set. This is only a heuristic measure with a performance increase bounded from above by some constant factor, and is sensitive to the number of splits and the distribution of exclusion windows. More splits also implies a quadratic growth in the number of grid-boxes and hence overhead. Still, we found this grid performed better in practice than using Python's *intervaltree* package once for each dimension, or the *r-tree* package. Faster implementations may be possible with alternative implementations of these data structures, segment trees or other principled methods of searching 2D space for point containment/rectangle overlap.

##### 6.2.2 Area Calculations

After locating all exclusion windows overlapping a given query RoI, Non-Overlap and Intensity Non-Overlap must then perform area calculations on them. Intensity Non-Overlap must separate all the subregions where a unique combination of exclusion windows overlaps the query RoI — Non-Overlap requires the area where only the query RoI is present and so is a special case of this more general procedure. Although these areas themselves are not rectangular, they form a disjoint collection of rectilinear polygons (informally, shapes whose sides can all be aligned with the axes) which can be covered by non-overlapping (axis-aligned) rectangles. By doing so, we can use only rectangles to describe the areas where only a particular combination of rectangles (e.g. RoIs/exclusion windows) overlap, despite these regions having more complicated shapes. In our case this allows us to compute the area of these regions just by summing the areas of these rectangles. (An example of an alternative algorithm would be to calculate the areas of all rectangular intersecting regions without subtracting any overlapping subregions, then update them starting from the region with the most intersections and working outwards.)

It is simple to decompose two overlapping rectangles into non-overlapping rectangles: the region in which they overlap is just a rectangle, and the 0-2 non-overlapping rectilinear polygons can be covered by 0-4 rectangles. More complicated situations arise when we consider the intersection of three or more boxes and the various combinations of their overlapping and non-overlapping regions. Then an obvious algorithm is to consider the boxes one-at-a-time, maintaining a set of non-overlapping boxes after every iteration. An algorithm of this sort allows updates fully online, but in our case the set of exclusion windows is only updated at the end of every injection. During injections, we need only consider query RoIs one-at-a-time without modifying the set of split exclusion windows — we only need to split the exclusion windows in batches between injections. Additionally, ideally we would like to cover the areas using the minimum number of rectangles necessary, but this simple recursive scheme will not in general produce output that is optimal in this sense.

To fit these requirements we use a form of line-sweep algorithm. For each rectangle, there are two endpoints on each dimension where the rectangle begins and ends (i.e. each rectangle is the cartesian product of two 1D intervals). We first sort all x-endpoints and then iterate through this ordered list. As rectangles begin, we store them as "active" rectangles, and as they end, we remove them from this storage. Whenever we would update the "active" rectangles, we can then sort their y-endpoints and iterate through them similarly. If we simply emit one rectangle for each adjacent pair of y-coordinates we iterate over this way, we will end up with a set of split non-overlapping rectangles — it is not possible for necessary splits to our rectangles to appear in locations where no original rectangle had an edge. However, this will lead to obvious cases where rectangles will be split unnecessarily. For example, if two rectangles intersected on the y-dimension but not the x-dimension, they would be partitioned into smaller rectangles whenever the other began or ended despite this being unnecessary.

Therefore we must do bounds-checking of active rectangle y-bounds, and in order to quickly query the active rectangles we store them in an interval tree. However, note that x-endpoints in this active tree will often partially overlap. When encountering a new incidence of this we can emit a box with height equal to the length of the entire interval, truncate said interval so it now only covers the non-overlapped length, and then create a new interval for the overlapping length. We therefore maintain *two* interval trees, which respectively contain the original active intervals and the split active intervals. Then when we receive a new endpoint, we can truncate all overlapping intervals in the split tree and emit a new rectangle for each, then repopulate the "missing" space using new intervals made by performing a y-sweep over the intervals returned by a query to the tree with the originals. In the geometric metaphor of the line-sweep, we can think of a vertical line sweeping along the x-endpoints. As it contacts them, a horizontal line performs a vertical sweep only along the length of the new endpoint, and after that, the vertical line "fills" the rectangle behind it. When endpoints only partially overlap, these fills split into two. By continuing this procedure to its end, we eventually fill in all the new rectangles i.e. obtain a set of non-overlapping rectangles.

##### 6.3 Worked Example

To illustrate the concept of Non-Overlap, let us take the (not necessarily realistic) example shown in Figures 14, 15 and 16. These figures show an example where we have three boxes $a$ (red), $b$ (blue) and $c$ (yellow). On the left is shown a breakdown of these three boxes into non-overlapping rectangular subregions, coloured to show their parents, and on the right is shown an intensity heatmap, where on a colourscale from grey to yellow to red each subregion is coloured according to the most intense parent box present. Figure 14 just shows this information for all boxes. Box $a$ can be seen to be least intense, and $c$ the most. Suppose we have previously fragmented some point in each of $a$ and $b$ previously (i.e. they are exclusion windows, and their intensity is their maximum fragmentation intensity) and we are interested in computing a score for whether we should fragment box $c$ (i.e. it is an active RoI, and its intensity is the intensity of its current precursor).

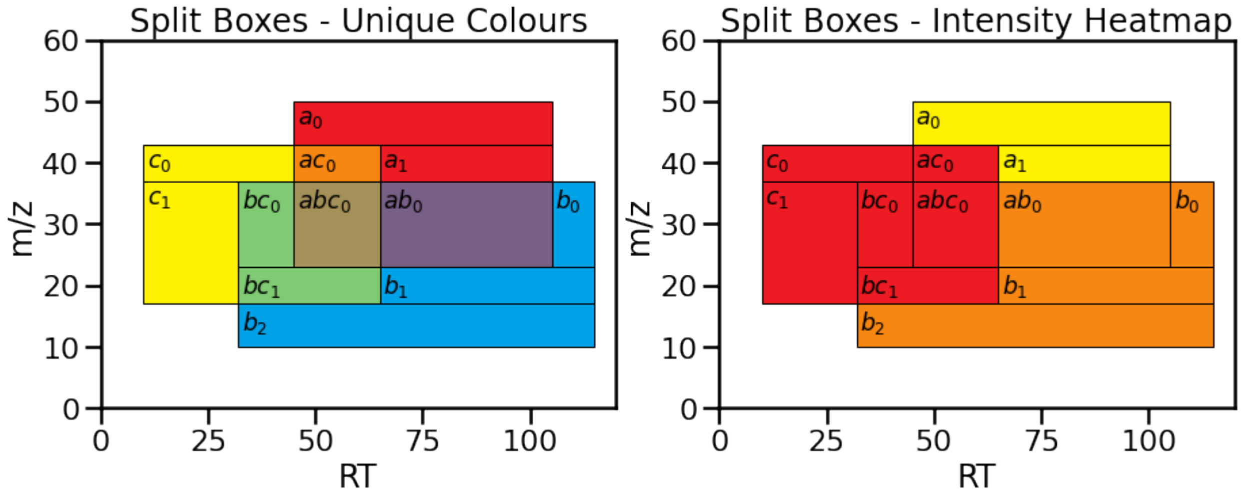


Figure 14: *Left:* A dummy example of how three overlapping boxes can be split into subregions, for illustration purposes. The three original boxes are coloured red, blue and yellow, and their shared areas interpolate their colours. *Right*: A heatmap of the intensities of the same boxes.

Then for the Non-Overlap score, we would use only the regions which are not covered by any other box (sum of all boxes $c_i$ whose names only contain $c$, e.g. $c_0, c_1, ...$) as a proportion of the total area (sum of all boxes $c_i^*$ with a name containing $c$, e.g. $c_0, c_1, ..., ac_0, ...bc_0, bc_1, ...abc_0, ...$). The boxes used for the numerator are highlighted in colour at their original intensity in Figure 15. Let $\lambda_{mod}$ be the log modified intensity, and all other symbols as in the main manuscript. Then in this case:

$$\lambda_{mod} = log(\lambda_c) \cdot \frac{\sum_i area(c_i)}{\sum_i area(c_i^*)}$$

$$\lambda_{mod} = log(\lambda_c) \cdot \frac{area(c_0) + area(c_1)}{area(c_0) + area(c_1) + area(ac_0) + area(bc_0) + area(bc_1) + area(abc_0)}$$

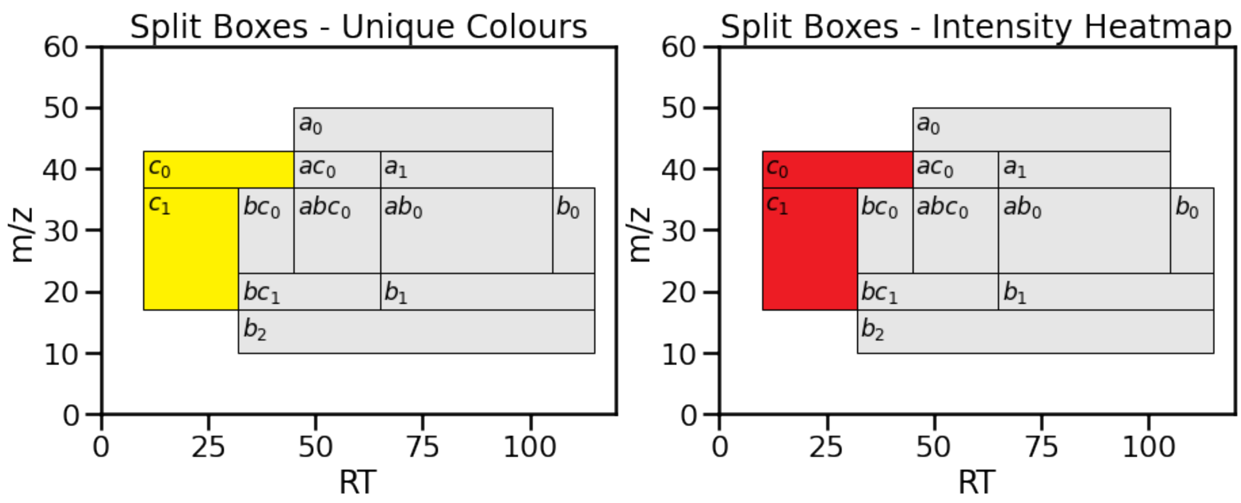


Figure 15: Example of how the Non-Overlap score would be calculated for $c$ (the leftmost, yellow box) in Figure 14. Anything unused for the calculation of the numerator is marked in grey. Non-Overlap exclusively uses the regions where only $c$ is present, and uses their intensities unmodified.

Having obtained the Non-Overlap modified intensity value from this, we need only apply our within-sample exclusion filters (DEW/SmartRoI/WeightedDEW) and intensity filter to get the final score. In addition to these values, Intensity Non-Overlap will also use the regions overlapped by $a$ and $b$, subtracting their intensities from the regions in $c$ (thus all boxes with $c$ in their label will be used in both the numerator and denominator). To assign intensities to these regions, for example, $\lambda_{cab}$ is the difference between the current precursor intensity of the RoI $c$ and the maximum fragmentation intensities of exclusion windows $a$ and $b$. This term may be negative (due to $a$ or $b$ being fragmented at a higher intensity) so we floor its contribution to the equation to zero. That the intensities of the overlapped regions are reduced can be seen in Figure 16. Now:

$$prop(c, B) = \frac{\sum_i area((cB)_i)}{\sum_i area(c_i^*)}$$

$$\lambda_{mod} = log\left(\sum_B max\left(0, \lambda_{cB}^{prop(c,B)}\right)\right)$$

$$\begin{aligned}
exp(\lambda_{mod}) = & \; max\left(0, pow\left(\lambda_c, \frac{area(c_0) + area(c_1)}{area(c)}\right)\right) \\
& + max\left(0, pow\left(\lambda_{ca}, \frac{area(ac_0)}{area(c)}\right)\right) \\
& + max\left(0, pow\left(\lambda_{cb}, \frac{area(bc_0) + area(bc_1)}{area(c)}\right)\right) \\
& + max\left(0, pow\left(\lambda_{cab}, \frac{area(abc_0)}{area(c)}\right)\right)
\end{aligned}$$

Once we apply the filters to this modified intensity we will get the final score.

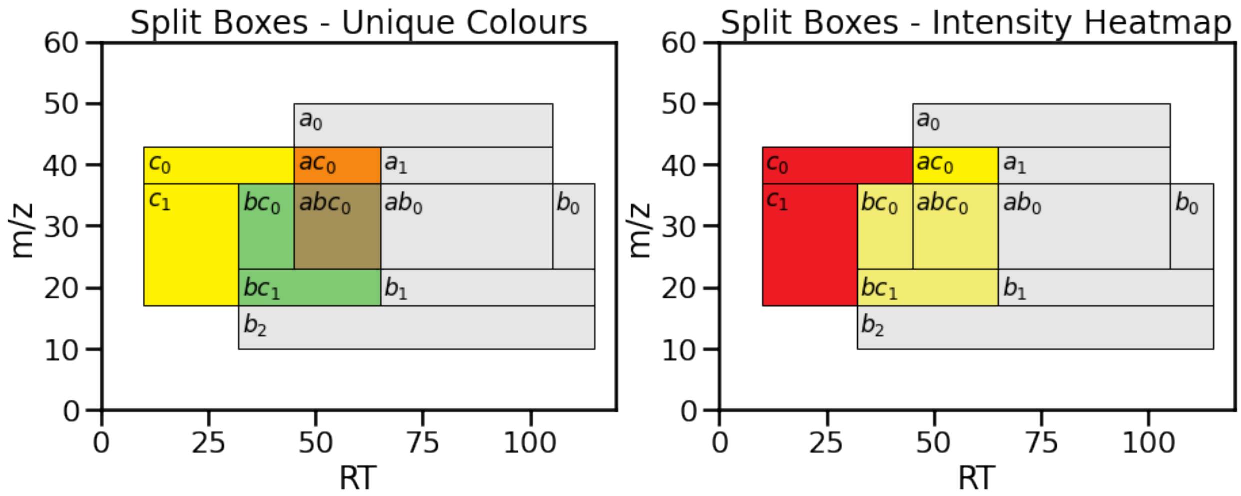


Figure 16: Example of how the Intensity Non-Overlap score would be calculated for $c$ (the leftmost, yellow box) in Figure 14. Anything unused for the calculation of the numerator is marked in grey. Note that all of the boxes touched by $c$ are used in the intensity area calculation. Also note that the overlapping areas used in Intensity Non-Overlap, but not Non-Overlap, are at decreased intensity compared to Figure 14 as they use the difference between $c$ (query RoI) and $a$ and $b$ (exclusion windows) where they are present.
